# Analysis of dRIF Composition Identifies ZNF346 as a Regulator of PKR Activation

**DOI:** 10.64898/2026.06.03.729416

**Authors:** Gaia Loucas, Roy Parker

## Abstract

Mammalian dsRNA sensors coordinate an array of cellular responses upon detection of dsRNA including translation repression, RNA degradation and interferon induction. Cytoplasmic endogenous or exogenous dsRNA can condense into cytosolic ribonucleoprotein assemblies in mammalian cells, referred to as dsRNA-induced foci (dRIFs), which enrich some dsRNA sensors including OAS3 and PKR. However, the protein composition of dRIFs, and their impact on dsRNA response regulation, are not well-understood. To determine other components of dRIFs, we reconstituted dRIFs in *in vitro* and analyzed their composition by mass spectrometry. We identified, and validated, multiple new dRIF components including dsRNA binding proteins and canonical RNA binding proteins with roles in innate immunity, transcription regulation, and proteostasis. Moreover, we identify dRIFs forming during mitosis with altered composition. Functional interrogation of dRIFs identified DHX9 as a limiting factor for dRIF formation, and the dRIF resident protein ZNF346 as a modulator of PKR signaling.

## INTRODUCTION

The cellular stress response to double-stranded RNA (dsRNA) is highly conserved, and has diverse downstream consequences for cell homeostasis (Y. G. Chen & Hur, 2022). dsRNA is a pathogen-associated molecular pattern (PAMP) and a hallmark sign of viral infection (Gantier & Williams, 2007). Subsequently, mammalian cells have evolved an array of distinct sensors to initiate the innate immune response to dsRNA (J. Wu & Chen, 2014). In addition, mounting evidence shows that sources of “self”-dsRNA including retroelement-derived and mitochondrial RNAs can also be activators of dsRNA sensors (J. Wu & Chen, 2014). While endogenous (“self”) dsRNA is generally sequestered in the mitochondrial and nuclear compartments of mammalian cells, mislocalization of “self”-dsRNA to the cytosol can trigger dsRNA sensors in the context of multiple chronic disease conditions (Y. G. Chen & Hur, 2022). These disease states are associated with sterile inflammation, cell cycle dysregulation, and dsRNA sensor hyperactivity, sometimes in cases where the activating ligand is unclear (Kim et al. 2019). Neurons may be particularly vulnerable to dsRNA accumulation, as long 3’ UTRs containing dsRNA regions are more abundant in brain tissue relative to other cell types, and may contribute toward inflammation and neuronal death observed in neurodegenerative disease (Dorrity, 2023).

The activation of different dsRNA sensors has multivariate cellular consequences. Upon detection of dsRNA, protein kinase R (PKR) phosphorylates eIF2*α* leading to translation arrest (Gal-Ben-Ari et al., 2018; Hartmann, 2017). OAS3 inhibits translation by synthesizing 2’-5’ oligoadenylate molecules in response to dsRNA, which activates RNase L and leads to global mRNA degradation (Hartmann, 2017; Y. Li et al., 2016). The dsRNA sensors RIG-I and MDA5 initiate a Type I Interferon (IFN-I) signaling cascade that promotes both proinflammatory (cell survival) and proapoptotic signaling (C.-P. Chan & Jin, 2022; Maelfait et al., 2020). Conversely, ADAR1 can dampen the immune response by disrupting dsRNA structures through its RNA editing properties (Liddicoat et al., 2015), thereby limiting dsRNA recognition by other dsRNA binding proteins (dsRBPs). Therefore, selectively modulating the activation of different dsRNA sensors can lead to different cellular fates in response to foreign or “self”-dsRNA-induced stress. However, the mechanisms that determine the outcome for an individual cell are obscured by the complexity of dsRNA signaling pathways.

We, and others, have observed ribonucleoprotein (RNP) granules that form due to the presence of cytoplasmic dsRNA in mammalian cells, which we refer to as dsRNA-induced foci (dRIFs) (Corbet et al., 2022) (Zappa et al., 2022). These assemblies enrich dsRNA and a number of dsRBPs, including PKR, ADAR1 (Corbet et al., 2022), and OAS3, the latter of which has been shown to recruit RNase L to dRIFs and enhance its ribonuclease activity (Cusic & Burke, 2024). dRIFs were originally described based on transfection of dsRNA into cells. However, dRIFs (i.e., cytoplasmic puncta of dsRNA, PKR, and other dsRBPs), are observed in multiple biological contexts. For example, dRIFs are observed in measles (Zappa et al., 2022), Dengue, and Zika infections in cultured cells (Cusic & Burke, 2024).

Additional observations suggest that dRIFs may also be relevant toward chronic disease states. Reversible PKR-enriched assemblies form during late mitosis (Zappa et al., 2022), which is consistent with data showing PKR activation and binding to inverted repeat Alu (IR-Alu) dsRNAs during cell division (Kim et al., 2014). These findings suggest that dRIFs may play a role in regulating cell cycle progression. Cytoplasmic dsRNA puncta have also been observed in multiple neurodegenerative conditions. Both dsRNA inclusions and p-PKR foci have been observed in the post-mortem brain tissue of ALS-FTD patients (Rodriguez, 2021). In addition, tau and TDP-43 loss-of-function phenotypes have been linked to dsRNA accumulation (Milstead, 2023). Whether formation of dsRNA foci in these conditions is a downstream biomarker of, or direct contributor to, neurodegenerative disease progression and etiology is an open question.

Previous work suggests that dRIFs are sites that promote PKR activation and subsequent phosphorylation of eIF2*α*, initiating translation arrest in a spatiotemporally regulated manner (Corbet et al., 2022). In addition, recent work indicates that enrichment of OAS3 and RNase L in dRIFs enhances OAS3 and RNase L activation upon dsRNA induction or viral infection (Cusic & Burke, 2024). Initial analyses of dRIFs suggested that specific dsRBPs are enriched in dRIFs whereas others are excluded. For instance, the dsRBPs RIG-I and MDA5 were not detected in dRIFs by immunofluorescence (Corbet et al., 2022). The selective enrichment of specific dsRBPs in dRIFs, and not others, may be related to whether their binding mechanism depends on cooperativity through homo-oligomerization, as is the case for IFN-I response-regulating sensors such as MDA5 (Berke et al., 2012; Peisley et al., 2011).

An important goal is to understand the molecular components of dRIFs, how those components influence dRIF assembly and persistence, and how that impacts the selectivity of the dsRNA response. The latter question is of particular interest, since single cell analyses have shown that individual cells within a single population can activate different axes of the dsRNA response (Burke et al., 2019). Given this observation, we developed a method to reconstitute and purify dRIFs from cell extracts and analyze their composition. In this work, we identify and validate new protein components of dRIFs by mass spectrometry. Analysis of enriched dRIF components identified the DExH-box helicase DHX9 as a component that limits dRIF formation, coupled with suppression of dsRNA response signaling. In addition, we identified the zinc finger protein ZNF346 as a dRIF resident protein and a previously unestablished activator of PKR.

## RESULTS

### dsRBP Granules Assembled in Cell Lysate (dRIF_IV_) Recapitulate *Bona Fide* dRIFs

To date, the comprehensive protein composition of dRIFs has not been confirmed. Therefore, our first goal was to obtain an assessment of the dRIF proteome. Initial attempts to purify dRIFs formed *in cellulo* were limited by the abundance of dRIFs in live cells. Given this, we adapted a previously developed *in vitro* reconstitution strategy that has been shown to faithfully recapitulate the compositional properties of stress granules and the granular component of nucleoli from lysed cells (Freibaum, 2021). For *in vitro* reconstitution of dRIFs, we used lysates from cells expressing mApple-PKR^K296R^, a catalytic mutant of PKR with higher binding affinity for dsRNA than wild-type PKR (Sharp, 1995), which we anticipated would increase dRIF stability. Our proposed approach for isolating dRIFs and analyzing their proteome is shown in Figure 1A.

**Figure 1.**
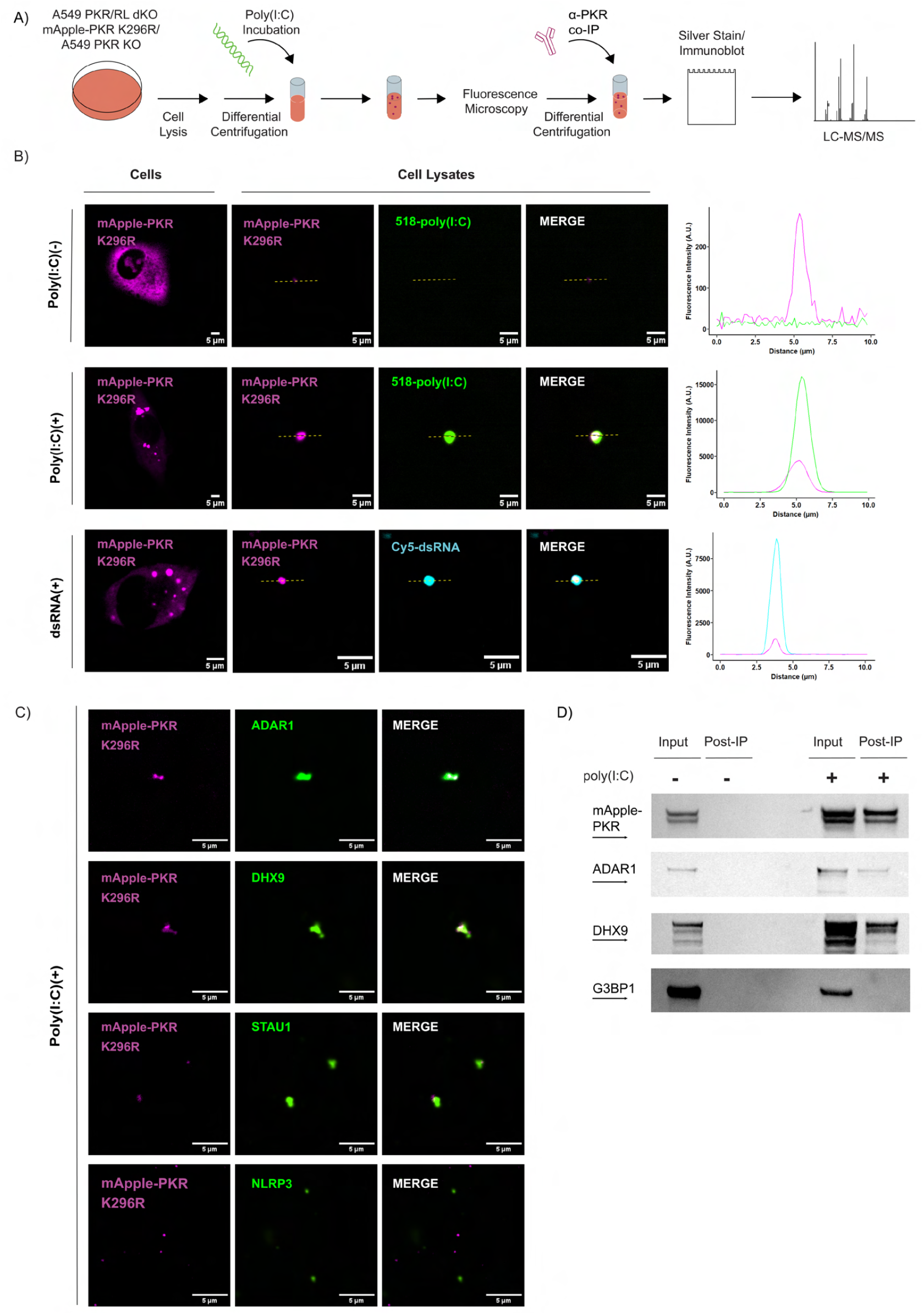
(**A**) Schematic for the formation, isolation, and co-IP of dRIFs reconstituted from cell lysates (dRIF_IV_) followed by analysis via MS to assess dRIF protein composition. (**B**) Fluorescence microscopy images of mApple-PKRK296R lysates with or without the addition of fluorescein-labeled poly(I:C) or cy5-labeled *in vitro*-transcribed dsRNA. Line scans are shown to assess overlap of mApple-PKR and poly(I:C) or dsRNA intensity profiles. Images of dRIFs formed in cells expressing mApple-PKRK296R are shown for reference. (**C**) Fluorescence microscopy images of cell lysates assessing colocalization between poly(I:C)-mApple-PKRK296R foci and antibody staining for other known dRIF resident proteins (ADAR1, DHX9, STAU1) compared to proteins not expected to enrich in dRIFs (NLRP3). (**D**) Western blot analysis assessing the enrichment of known dRIF constituents (mApple-PKR, ADAR1, DHX9) and the exclusion of proteins known to form independent granules (G3BP1) in immunopurified dRIF_IV_.

Our first task was to assess the viability our *in vitro* reconstitution approach to forming dRIFs. We treated cell lysates with fluorescein-labeled poly(I:C) and inspected them for the formation of dsRNA-dsRBP assemblies, which we assessed by looking for the formation of mApple-PKR- and fluorescein-positive foci via fluorescence microscopy. As anticipated, we observed spontaneous formation of poly(I:C)-PKR co-assemblies within 30 minutes of poly(I:C) addition, or of *in vitro* transcribed dsRNA (Figure 1B). This demonstrates that dsRNA assemblies containing PKR can form in cell lysates, which we refer to as dRIF_IV_, with the subscript IV to denote assembled *in vitro*.

To determine if dRIF_IV_ were compositionally similar to *bona fide* dRIFs, we immuno-stained dRIF_IV_ for proteins known to concentrated or be excluded from dRIFs in cells. We observed that the dRIF components ADAR1, DHX9, STAU1 also colocalized with dRIF_IV_ (Figure 1C). In contrast, the NLRP3 protein, which is not enriched in dRIFs in cells (Corbet et al., 2022), did not co-assemble into dRIF_IV_ (Figure 1C). This demonstrates that assemblies of dsRNA, PKR and other dsRBPs can form in cell lysates, which resemble physiological dRIFs.

It was imperative to determine whether dRIF_IV_ were sufficiently stable for isolation and downstream analysis. To this end, we performed differential centrifugation to enrich large molecular assemblies, followed by affinity purification with an *α*-PKR antibody. These assemblies persisted through the co-immunoprecipitation (co-IP) process, as indicated by microscopy analysis of the beads 24 hours after the initial step of poly(I:C) induction (Figure S1A).

To confirm whether dRIF_IV_ retained other components of dRIFs throughout the purification process, we performed immunoblotting for several proteins previously demonstrated to localize to dRIFs (Corbet et al., 2022). We observed the previously identified dRIF proteins (PKR, ADAR1, STAU1) enriched in dRIF_IV_ (Figure 1D). In contrast, G3BP1, which is absent from dRIFs *in cellulo* and is a key component of stress granules, was excluded (Figure 1D). These observations argue that dRIF_IV_ reflect many of the compositional properties of dRIFs formed in live cells.

Consistent with dRIF-like assemblies forming in cell lysates, the addition of either poly(I:C) or dsRNA was required for the formation of these assemblies and the co-IP of known dRIF components with PKR (Figure 1A, 1D). Furthermore, cell lysates from a PKR KO cell line that were treated with poly(I:C) and immunopurified using a *α*-PKR antibody did not show visible enrichment of dRIF proteins by western blot analysis (Figure S1B). These observations indicate that our *in vitro* reconstitution and co-IP strategy isolates assembles resembling physiological dRIFs.

### Mass Spectrometric Analysis of dRIF_IV_ Composition

After isolating dRIF_IV_ from cell lysates by subcellular fractionation and *α*-PKR co-IP, we submitted these samples for analysis by label-free quantitative mass spectrometry (LFQ-MS). Analysis of the MS results identified 35 proteins that were enriched in dRIF_IV_ as compared to controls where PKR KO cells were processed in parallel using the same strategy, or where PKR was immunopurified without the addition of dsRNA (Figure 2A-B). Proteomics analysis of these preparations identified two broad classes of factors enriched in dRIF_IV_.

**Figure 2.**
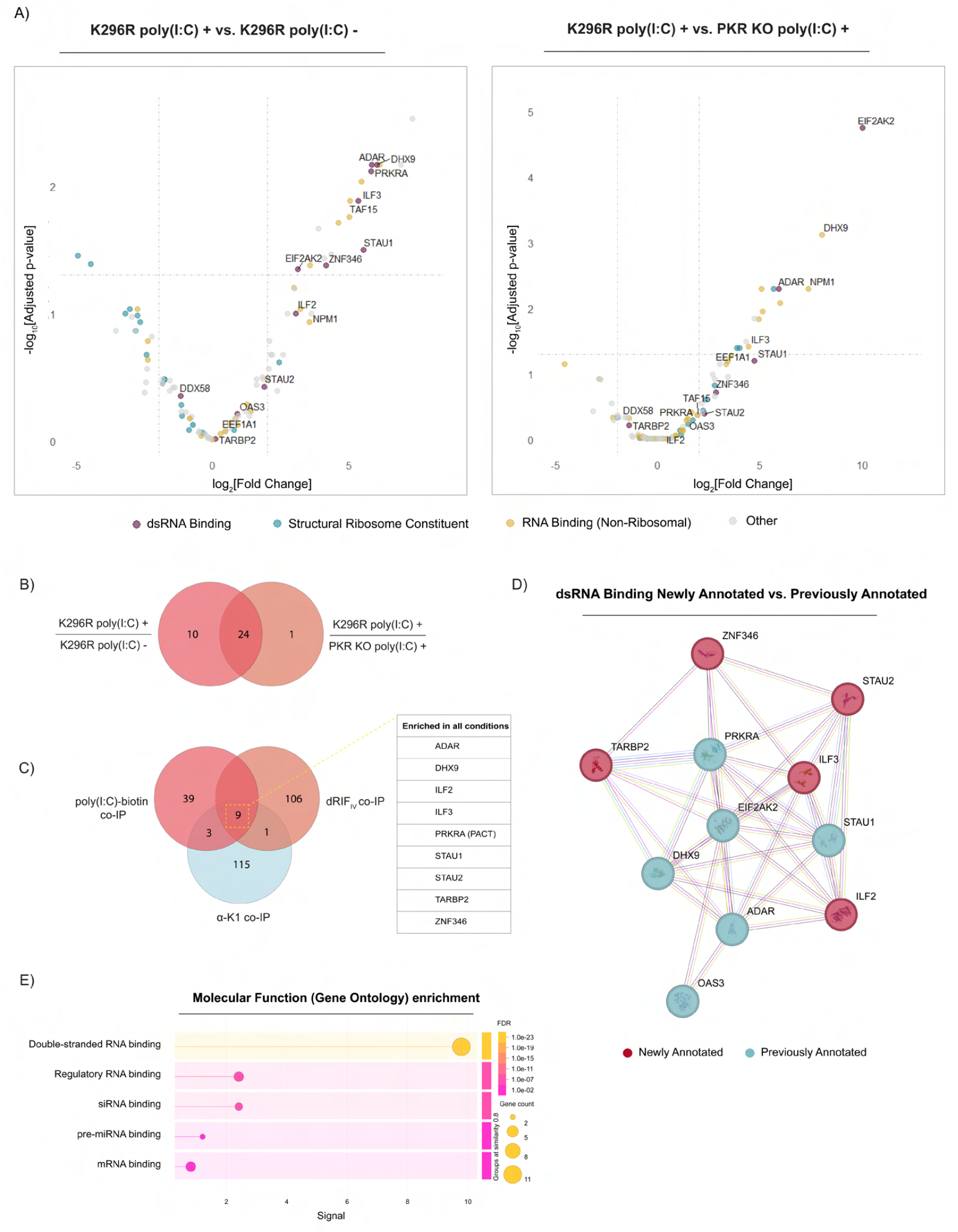
(**A**) Results from MS analysis of dRIF_IV_. Volcano plots represent differentially enriched proteins against no-poly(I:C) controls (left) and lysates from a PKR KO cell line that were treated in parallel with poly(I:C) (right). The log2[Fold Change] is calculated as the difference between four technical replicates in the (K296R poly(I:C)+ group) and two replicates each of the (K296R poly(I:C)-) or (PKR KO poly(I:C)+) negative controls. Replicates were obtained from 2 independent co-IP assays. (**B**) Summary of detected proteins that were enriched (log2[Fold Change] value greater than 1 and a false discovery rate of less than 0.05) compared to one or both negative control samples (lysates without poly(I:C) added or PKR KO cells with poly(I:C) added). (**C**) Comparison of overlapping proteins from dRIF_IV_ co-IP with proteins identified by Lim et al. (2025) using either biotinylated poly(I:C) or *α*-K1 antibody as baits. (**D**) STRING interactome depicting dsRBPs detected by MS previously reported to enrich in dRIFs (blue), and newly annotated dRIF candidate dsRBPs (pink). (**E**) STRING database plot showing the Gene Ontology (GO) Molecular Function terms for the enriched dsRBPs shown in C).

First, an array of dsRBPs were identified by MS (Figure 2A,E). Some of these were anticipated since they are previously known dRIF constituents (EIF2AK2 (PKR), ADAR1, STAU1, DHX9, PACT, OAS3). In addition, we identified several dsRBPs as new putative dRIF components including ILF3, STAU2, and ZNF346 (Figure 2D). In agreement with previous findings (Corbet et al., 2022), the MAVS-activating dsRNA sensor MDA5 was absent from these preparations. This may be due to the requirement of MDA5 oligomerization along tracts of dsRNA for stable binding (B. Wu et al., 2013). The other MAVS-activating dsRNA sensor RIG-I (DDX58) was detected but its enrichment with PKR co-IP was decreased relative to other dsRNA sensors, supporting the interpretation that dRIF_IV_ possess a degree of compositional selectivity for select dsRNA sensors and other dsRBPs.

Second, we identified several proteins by MS in dRIF_IV_ that are not canonically associated with dsRNA binding, but are known to bind single-stranded RNA (ssRNA) (Figure 2A). ssRNA-binding proteins enriched in dRIF_IV_ by MS included ANXA2, ILF2, TAF15, NPM1, and EEF1A1 (Figure 2A). We were encouraged to find that many of these ssRNA-binding proteins were non-ribosomal (Figure 2A), as ribosome constituents are common background contaminants in mass spectrometry (Mellacheruvu, 2013). Some factors enriched in dRIF_IV_ are known to affect dsRNA metabolism, either through remodeling or stabilization of dsRNA. ADAR1 resolves dsRNA duplexes through its A-I editing activities, and is subsequently associated with immunosuppression during viral infection (Pfaller et al., 2018). In addition, recent work has revealed DHX9 and ADAR1 as critical suppressors of immune responses to endogenous dsRNAs (Cottrell et al., 2024; Liddicoat et al., 2015; Zhou et al., 2024). ILF3 binds viral RNAs to inhibit their replication (Harashima et al., 2010), and modulates PKR by regulating the synthesis and trafficking of PKR-inhibiting circRNAs (C.-X. Liu et al., 2019). Moreover, ILF2/3 are reported to form heterodimers (Wolkowicz & Cook, 2012) and antagonize Staufen-mediated decay by binding to structured 3’ UTRs targeted for degradation by STAU1/2 (Nourreddine et al., 2020). The enrichment of dsRBPs that modulate dsRNA stability and immunogenicity in dRIF_IV_ supports a role for these assemblies in dsRNA response regulation.

### Comparison of dRIF_IV_ Components to the dsRNA Interactome

To validate our dRIF_IV_ proteome, we first compared our findings to recent work that analyzed the dsRNA interactome by mass spectrometry (Lim et al., 2025). Many of the proteins from our proteomics analysis of dRIF_IV_ were in agreement with this interactome analysis, which identified dsRBPs through co-IP with dsRNA using either biotinylated poly(I:C) or an *α*-K1 (*α*-dsRNA) antibody (Lim et al., 2025). In total, 12 of the proteins identified in our analysis overlapped with the biotinylated poly(I:C) bait and 10 overlapped with the K1 bait (Figure 2C). Of these proteins, 9 of them were enriched by all 3 co-IP methods (Figure 2C). This indicates that at least 9 of the proteins enriched in dRIF_IV_ can bind endogenous dsRNAs, and are not artifacts of poly(I:C) treatment. Moreover, since the samples prepared by K1 co-IP were treated with ssRNase (Lim et al., 2025), we concluded that the 9 overlapping proteins were recruited either in a dsRNA-dependent manner, or indirectly through protein-protein interactions with dsRBPs. The latter possibility likely applies to ILF2, which lacks dsRNA binding activity but interacts with ILF3 independently of its dsRBDs (Bonczek et al., 2022).The only other overlapping protein that lacked canonical dsRBDs was NPM1. Despite this, NPM1 exhibits enhanced dsRNA binding after IFN*γ* and TNF*α* stimulation (Pang et al., 2003), further implicating NPM1 in dsRNA regulation. Thus, comparison of dRIF_IV_ to these dsRNA/poly(I:C) interactomes argues that our purification method identified *bona fide* dRIF components.

### Validation of dRIF Constituents

A key question is whether new proteins identified in dRIF_IV_ by mass spectrometry are also present in cellular dRIFs. Given this, we used fluorescence microscopy *in situ* to examine if a subset of proteins identified in dRIF_IV_ were also enriched in cellular dRIFs. Most of these proteins were selected based on agreement with the dsRNA interactome (Figure 2C). We visualized candidate dRIF proteins by either immunofluorescence (IF) staining, and/or by expression of a fluorescently tagged derivative of the protein of interest in the A549 mApple-PKR^K296R^ cell line used for dRIF_IV_ isolation. To ensure that identified proteins were recruited to dRIFs in a dsRNA-dependent manner, and were not specific to poly(I:C), sequence-specific dsRNA was transfected into cells for validation of dRIF components.

*In situ* analysis confirmed that several proteins enriched in dRIF_IV_ were *bona fide* dRIF components in cells. First, by IF we observed that both the dsRBP ILF3 and its heterodimeric partner ILF2 localized to dRIFs induced by exogenous *in vitro* transcribed dsRNA (Figure 3A-3B), with ILF3 colocalizing with mApple-PKR foci 35% of the time and ILF2 colocalizing 23% of the time (Figure 3A-B). Second, NPM1, an RNA-binding protein enriched in our proteomics data set, is recruited to 23% of dRIFs in non-mitotic cells (Figure 3C). Third, IF staining showed that the translation elongation and transcription factor EEF1A1 localizes to 15% of dRIFs induced by exogenous dsRNA (Figure 3D). In addition, p62, which was present in some of the mass spectrometry analyses and is a major adaptor protein for both autophagy and proteolysis, colocalized with dRIF assemblies *in situ* 7% of the time (Figure 3E). Prior work has shown that p62 promotes the clearance of DHX9 stress granules, which can contain dsRNA (Zhou et al., 2024). This suggests the possibility that p62 recruitment to dRIFs may facilitate their clearance through autophagy.

**Figure 3.**
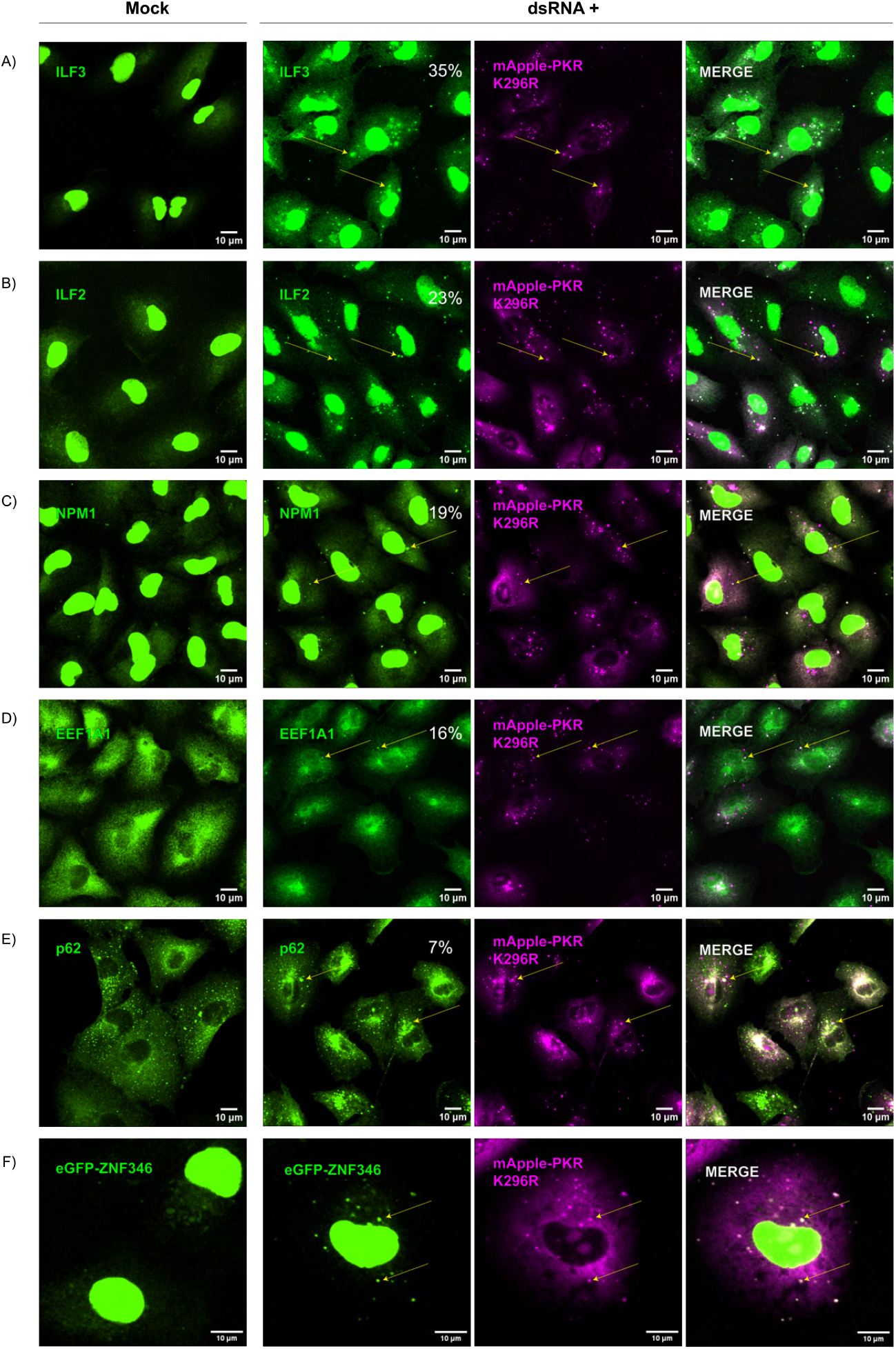
(**A-F**) Representative fluorescence microscopy images for *in situ* validation of dRIF candidate proteins identified by MS. Validation of candidates was performed by either immunostaining for the target protein or by expression of eGFP-tagged derivative of the target protein in A549 mApple-PKR^K296R^ cells, as indicated in the figure labels. Percentages indicate the frequency with which dRIF candidate proteins colocalized with mApple-PKR^K296R^ foci by fluorescence microscopy. For validation experiments, cells were transfected with *in vitro*-transcribed dsRNA to confirm that recruitment of candidate proteins to dRIFs is stimulated by dsRNA and is not specific to poly(I:C).

Due to poor antibodies for establishing ZNF346 localization by IF, we tested the recruitment of ZNF346 to dRIFs using an eGFP-ZNF346 construct. We observed that eGFP-ZNF346 co-localized with dRIFs following dsRNA transfection (Figure 3F). Interestingly, ZNF346 contains 4 zinc finger domains with high avidity for dsRNA which are required for its nucleolar localization (Yang et al., 1999). However, ZNF346 can be exported to the cytosol with ILF3 (T. Chen et al., 2004), and has been reported to co-IP with PKR (Buljan et al., 2020; Cho et al., 2022; S. Li et al., 2011; Wang et al., 2023) and multiple forms of dsRNA (Lim et al., 2025), Although the function of ZNF346, and how its localization relates to its function is not well-studied, these reports support the physiological relevance of eGFP-ZNF346 recruitment to dRIFs.

Despite its high enrichment in our proteomics data, we did not detect the E3 ligase TRIM21 in dRIFs by either IF or expression of a TRIM21-eGFP fusion protein, at either acute (3-4 hours) or longer (24 hours) timepoints (data not shown). Since TRIM21 scores moderately in a database of proteomic contaminants (Mellacheruvu, 2013), and is also a known IgG receptor (Keeble et al., 2008), we concluded that TRIM21 enrichment was an artifact of our proteomics pipeline. Similarly, enrichment of PCBP2 in dRIFs was not observed by immunostaining after transfection of sequence-specific dsRNA (data not shown). Since PCBP2 is a well-established binder of poly(C) oligomers, we interpret its absence in dRIFs *in situ* to the implementation of poly(I:C) for our dRIF isolation protocol.

Taken together, these observations demonstrate that many, but not all, of the components found in dRIF_IV_ enrich in dRIFs formed in cells. Interestingly, the observation that dRIF components are not always detected in each assembly in both this work (Figure 3) and earlier work (Corbett et al, 2022) highlights that individual dRIFs can differ in composition (see discussion).

### dRIFs Form During Mitosis

Prior results have identified PKR foci that form during mitosis (Zappa et al., 2022). These puncta form during mitosis independently of any exogenous dsRNA stimulus, and are presumably formed due to nucleation by endogenous IR-Alu dsRNAs released into the cytosol during mitosis (Kim et al., 2014). Since these assemblies would be predicted to be compositionally similar to dRIFs, we used fluorescence microscopy to examine whether proteins observed in dRIFs forming in non-mitotic cells were also present in mitotic PKR-enriched assemblies. This analysis revealed the following observations.

First, we observed that NPM1 frequently localized to PKR-containing puncta during late mitosis that persist through early interphase (Figure 4A), which we refer to as mitotic dRIFs. NPM1 has been suggested to inhibit PKR activity (Pang et al., 2003), and PKR activation has been shown to play a role in faithful mitotic progression (Kim et al., 2014). This suggests that enrichment of both NPM1 and PKR in mitotic “dRIFs” (Figure 4A) may promote mitotic exit by dampening PKR activation.

**Figure 4.**
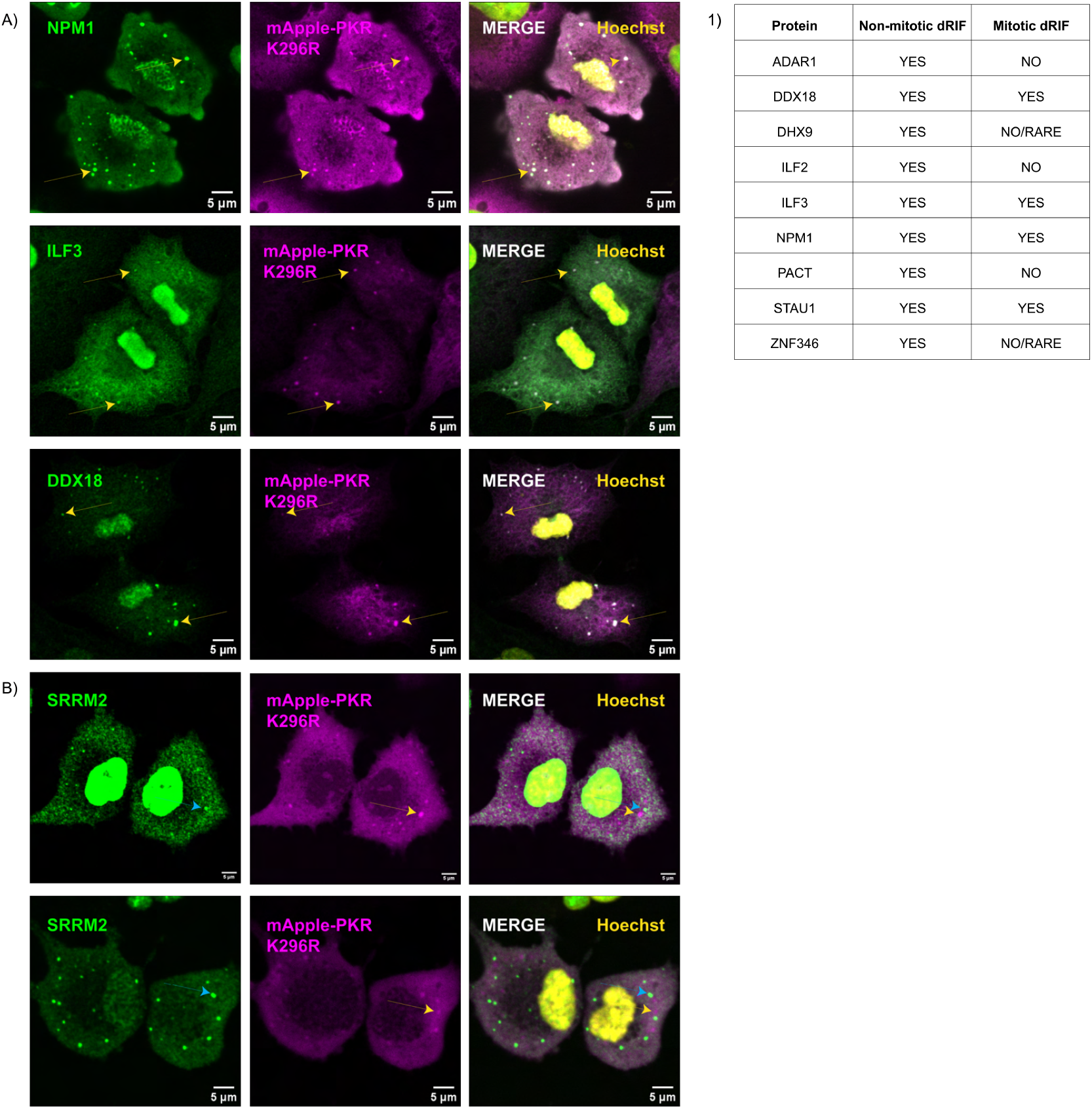
(**A**) Immunofluorescence microscopy images of select proteins (ILF3, NPM1, DDX18) which are enriched in exogenously induced dRIFs that are also enriched in mitotic/early interphase assemblies in the absence of exogenous dsRNA stimulus (“mitotic dRIFs”). (**B**) Immunofluorescence microscopy images showing that mitotic dRIFs (yellow arrows) are spatially distinct from MIGs (blue arrows), which are marked by puncta enriched with the splicing factor SRRM2. (**1**) Table comparing enrichment of proteins identified in dRIF_IV_ in exogenously induced dRIFs versus mitotic dRIFs.

Second, we observed that some proteins found in exogenously induced dRIFs are also concentrated into mitotic dRIFs, including ILF3 and STAU1 (Figure S4A). However, many dsRBPs seen in cytoplasmic dRIFs either enriched in mitotic dRIFs rarely or not at all. ADAR1 formed its own distinct puncta during late mitosis/early interphase that appeared to dock with mitotic dRIFs (Figure S4A). Similarly, ZNF346 formed assemblies during mitosis that did not colocalize with NPM1-enriched foci (Figure S4A). We only observed enrichment of DHX9 in a single mitotic dRIF within the 12 pairs of dividing cells we measured (Figure S4A). We did not observe any enrichment of ILF2 or PACT in mitotic dRIFs (Figure S4A). Since ILF3 and STAU1 compete for the same 3’ UTR structures in endogenous RNAs (Nourreddine et al., 2020), their preferential enrichment (Figure S4A) supports the possibility that specific endogenous dsRNAs are concentrated in mitotic dRIFs.

Both NPM1 and ILF3 localize to the nucleolus (W. Y. Chan et al., 1989; Viranaicken et al., 2011), and STAU1 can undergo nucleolar shuttling (Martel et al., 2006). Therefore, we wondered whether other nucleolar components were also enriched in mitotic dRIFs. Indeed, the RNA helicase DDX18, a key regulator of nucleolar condensation in coordination with NPM1 (X. Shi et al., 2024), was concentrated in some mitotic dRIFs by IF (Figure 4). We then immuno-stained dsRNA-treated cells and confirmed by that DDX18 is also a component of exogenously induced dRIFs (Figure S4B). Together, these observations show that mitotic dRIFs share components of dRIFs induced from exogenous, cytoplasmic dsRNA, but also possess enhanced concentrations of nucleolar constituents including NPM1 and DDX18.

### Mitotic dRIFs are Distinct from Mitotic Interchromatin Granules

Prior reports have described a type of mitotic ribonucleoprotein (RNP) granule, referred to as a mitotic interchromatin granule (MIG), that contains many nuclear proteins and snRNAs including the splicing factor SRRM2 (Lester et al., 2023; Prasanth et al., 2003; Rai et al., 2018). To determine if MIGs and mitotic dRIFs were overlapping or compositionally distinct RNP granules, we imaged cells for SRRM2 as a marker of MIGs (Lester et al., 2023) and mApple-PKR as a marker of mitotic dRIFs. We observed that MIGs and mitotic dRIFs form independent, spatially distinct foci (Figure 4B). Moreover, dRIFs are also distinct from cytoplasmic speckles (Figure 4D), which are MIG-like assemblies that form in non-mitotic cells (Lester et al., 2023). This demonstrates that dRIFs, both in mitosis and in non-mitotic cells, are distinct from SRRM2-containing assemblies.

### DHX9 Limits dRIF Formation and dsRNA Responses

While PKR and OAS3, which are both enriched in dRIF_IV_ and cytoplasmic dRIFs *in cellulo*, have been established as a key dRIF constituents, previous work has shown that neither are required for dRIF formation (Corbet et al., 2022; Cusic & Burke, 2024). Most of the dsRBPs identified in dRIF_IV_ by mass spectrometry have multivalent binding properties, i.e. they possess multiple dsRNA binding domains (Jeon & Kim, 2025). Moreover, some (ADAR1, PACT) are also reported to interface with other dsRBPs through their dsRBDs (Chukwurah et al., 2021; Sinigaglia et al., 2024) or through independent protein-protein binding motifs (Sanchez David et al., 2019). By analogy to other known RNP granules, these proteins are predicted to form multiplexed interactions with multiple molecules of (ds)RNAs and other proteins in dRIFs. Therefore, we anticipated that multivalent dsRBPs may serve as condensers of dsRNA, similar to the role of G3BP1/2 in condensing mRNAs to form stress granules (Kedersha et al., 2016; P. Yang et al., 2020). In addition, we expected RNA helicase activity to play a role in the disassembly of dRIFs, by analogy to RNA helicases that limit stress granule formation (Tauber et al., 2020).

We hypothesized that multivalent proteins that were enriched in dRIF_IV_ by MS were preferentially recruited to dRIFs based on 1) their multivalent properties and 2) their general cellular abundance. Five of these proteins (DHX9, ILF3, PACT, STAU1, ZNF346) were selected for experiments based on their multivalent properties and/or known roles in dsRNA regulation. We then sought to determine whether altering expression of these proteins could modulate dRIF formation or disassembly, and/or impact the dsRNA response.

To first determine whether any of the 5 selected dRIF constituents contributed toward dRIF formation, we performed siRNA-mediated knockdown of the target proteins and compared dRIF formation after dsRNA transfection in each knockdown condition (Figure 5A). We used an A549 PKR KO cell line expressing an mApple-PKR^K296R^ transgene for these experiments, since the reduced off-rate of this PKR mutant from dsRNA should increase the dynamic range of our visual readouts of dRIF formation. The presence of discrete cytosolic mApple-PKR^K296R^ puncta was used as the primary marker of dRIFs, supplemented by transfection of fluorescently labeled dsRNA as a secondary readout of dRIF formation.

**Figure 5.**
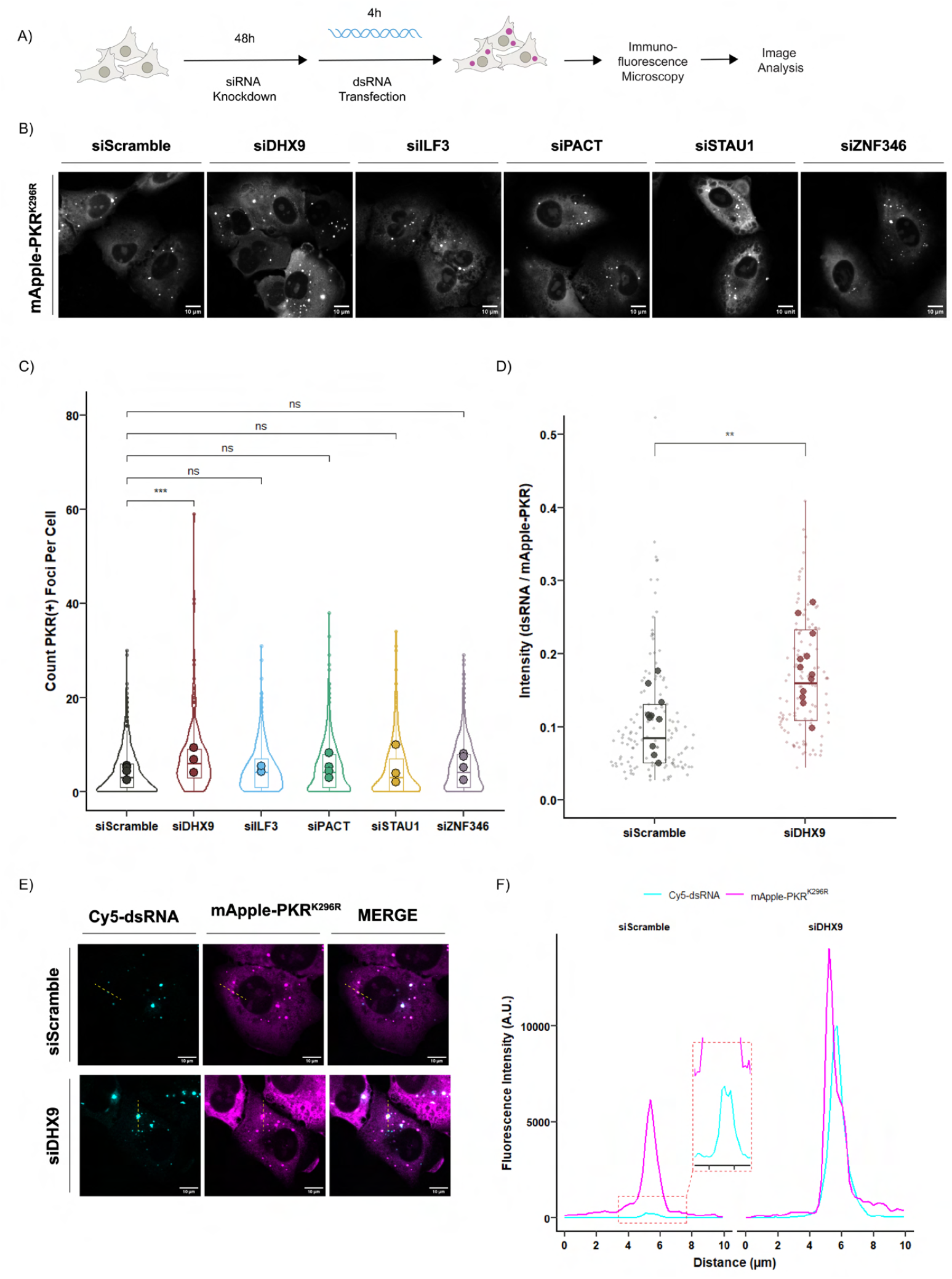
(**A**) Schematic of siRNA knockdown strategy to test the impact of depleting dRIF resident proteins on dRIF formation. A549 PKR KO mApple-PKR^K296R^ cells were treated with siRNA for 48 hours prior to dsRNA transfection. siRNA-treated cells were transfected with dsRNA for 4 hours and fixed for analysis by fluorescence microscopy. siRNA-treated cells were harvested in parallel for western blotting to assess knockdown efficiency. (**B**) Representative fluorescence microscopy images of cells with dRIFs (proxied by mApple-PKR^K296R^ foci) in each siRNA knockdown condition. (**C**) Quantification showing the number of dRIFs per cell, denoted by mApple-PKR^K296R^-positive foci, in each siRNA knockdown condition. The mean values for each biological replicate (calculated as the mean of per-image means for each rep) are shown as circles for each condition. Statistical comparisons were made using an unpaired two-tailed t-test on the per-image means for each siRNA knockdown condition across biological replicates. Data was obtained from 3-5 independent experiments per condition, 3+ images per experiment, and 50+ cells per biological replicate per condition. (**D**) Quantification of cy5-labeled dsRNA fluorescence intensity (normalized to mApple-PKR^K296R^ fluorescence intensity) in cells treated with siRNA targeting DHX9 (siDHX9) or a non-targeting control (siScramble). The per-image means are shown as larger circles, smaller circles represent individual data points. Statistical comparisons were made using a Wilcoxon rank-sum test on per-image means. Data was obtained from 2 independent experiments and 40+ cells and 3+ fields-of-view per condition and biological replicate. (**E**) Microscopy images measuring dsRNA fluorescence intensity (normalized to mApple-PKR^K296R^ fluorescence intensity) in cells treated with siRNA targeting DHX9 (siDHX9) or a non-targeting control (siScramble). (**F**) Fluorescence intensity profiles from the lines in 5E) showing the relative enrichment of cy5-dsRNA and mApple-PKR^K296R^ in dRIFs in single cells treated with siRNA targeting DHX9 (siDHX9) or a non-targeting control (siScramble). All data in this figure shows cells transfected with 400 ng/mL dsRNA. All statistical analyses in this figure use the following p-value cutoffs: ^****^P ≤ 0.0001, ^***^P ≤ 0.001, ^**^P ≤ 0.01, ^*^P ≤ 0.05, n.s. P *>* 0.05.

We observed that knockdown of DHX9 enhanced dRIF formation by several criteria. DHX9 knockdown both increased the number of dRIF-positive cells and increased the number of dRIFs in individual cells (Figure 5B-C). Moreover, DHX9-depleted cells had higher concentrations of dsRNA enriched in dRIFs relative to mApple-PKR^K296R^ (Figure 5D-F). The accumulation of dsRNA in dRIFs upon DHX9 depletion supports the interpretation that DHX9 mediates the clearance of dsRNA from dRIFs. In contrast, most of the other dRIF components tested did not significantly impact dRIF formation (Figure 5B-C), suggesting that dRIF assembly may be redundantly regulated by multiple dsRBPs.

Due to the compositional selectivity of dRIFs, we predicted that altering the levels of dRIF constituents could change the relative amplification of specific dsRNA response pathways. We therefore examined how knockdown of dRIF resident proteins impacted signaling along three distinct axes of the dsRNA response: MAVS-mediated activation of the IFN-I program, OAS3-RNase L-mediated activation of host mRNA decay, and PKR-mediated translation arrest (Figure 6A). We confirmed that our readouts for these responses increased in a manner dependent on dsRNA treatment in preliminary experiments (Figure S5C-E).

**Figure 6.**
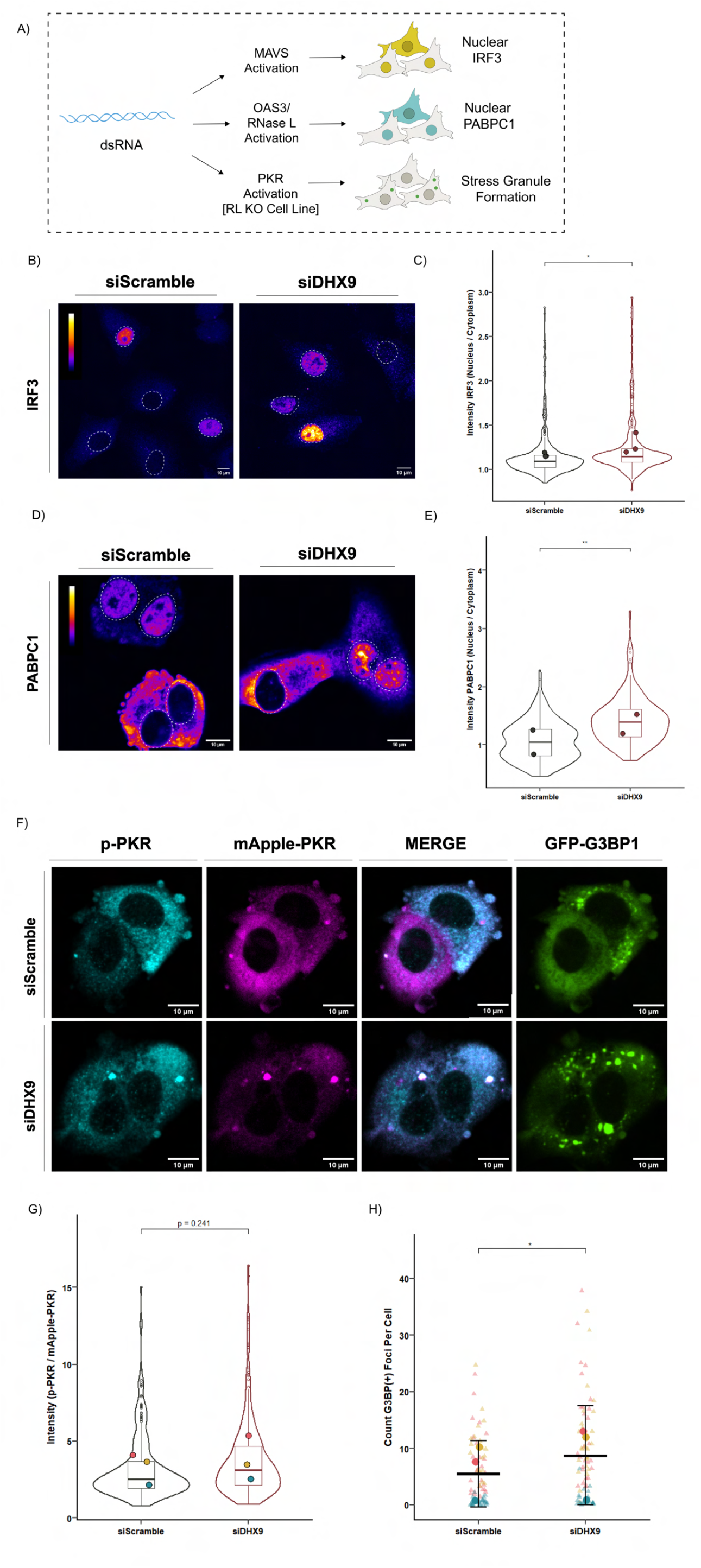
(**A**) Schematic of immunofluorescence assays for measuring IRF3 nuclear/cytoplasmic intensity to approximate MAVS activation, PABPC1 nuclear/cytosolic intensity to measure RNase L activation, and visualization of stress granules (denoted by GFP-G3BP1 in RNase L KO cells) as a marker of PKR-dependent translation arrest. (**B**) Representative fluorescence microscopy images showing the frequency of MAVS activation (marked by nuclear IRF3) in dsRNA-transfected cells treated with siRNA targeting DHX9 (siDHX9) or a non-targeting control (siScramble). Dashed lines represent nuclear borders. Intensity map represents relative gray scale intensity. (**C**) Quantification comparing the intensity ratios of nuclear to cytosolic IRF3 in dsRNA-transfected cells treated with siRNA targeting DHX9 (siDHX9) or a non-targeting control (siScramble). Statistical comparisons were made using a Wilcoxon rank-sum test on per-image means. Data was obtained from 3 independent experiments and 30+ cells and 2+ fields-of-view per biological replicate and condition. (**D**) Representative fluorescence microscopy images showing the frequency of RNase L-activation (marked by nuclear PABPC1) in dsRNA-transfected cells treated with siRNA targeting DHX9 (siDHX9) or a non-targeting control (siScramble). Dashed lines represent nuclear borders. Intensity map represents relative gray scale intensity. (**E**) Quantification showing the fluorescence intensity ratios of nuclear to cytosolic PABPC1 in dsRNA-transfected cells treated with siRNA targeting DHX9 (siDHX9) or a non-targeting control (siScramble). Statistical comparisons were made using a Wilcoxon rank-sum test on per-image means. Data was obtained from 2 independent experiments and 50+ cells and 3+ fields-of-view per biological replicate and condition. (**F**) Representative fluorescence microscopy images showing the relationship between PKR activation, dRIF formation (denoted by mApple-PKR foci) and the presence of stress granules (denoted by GFP-G3BP1 foci) in dsRNA-transfected RNase L KO cells treated with siRNA targeting DHX9 (siDHX9) or a non-targeting control (siScramble). (**G**) Quantification showing the mean intensity of p-PKR staining (normalized to mApple-PKR signal) in dsRNA-transfected RNase L KO cells treated with siRNA targeting DHX9 (siDHX9) or a non-targeting control. Statistical comparisons were made by performing an unpaired two-tailed t-test on per-image means. Data was obtained from 3 independent experiments, and 60+ cells and 5+ fields-of-view per biological replicate and condition. (**H**) Quantification showing the percentage of stress granule-positive cells (proxied by GFP-G3BP1-positive foci) in dsRNA-transfected RNase L KO cells treated with siRNA targeting DHX9 (siDHX9) or a non-targeting control. Statistical comparisons were made using an unpaired two-tailed t-test on per-image means. Data was obtained from 3 independent experiments, and 100+ cells and 5+ fields-of-view per biological replicate and condition. All data in this figure shows cells transfected with 400 ng/mL dsRNA. All statistical analyses in this figure use the following p-value cutoffs: ^****^P ≤ 0.0001, ^***^P ≤ 0.001, ^**^P ≤ 0.01, ^*^P ≤ 0.05, n.s. P *>* 0.05.

Consistent with DHX9 expression reducing dRIFs and their dsRNA enrichment, we observed that depletion of DHX9 enhanced the three measured cellular responses to dsRNA. First, we observed that knockdown of DHX9 led to increased nuclear translocation of IRF3 (Figure 6B-C), which occurs following activation of MAVS signaling by MDA5 or RIG-I binding dsRNA (S. Liu et al., 2015; Nabeel-Shah et al., 2022). Second, we observed that DHX9 knockdown increased the population of cells with activated RNase L as assessed by nuclear PABPC1 translocation (Figure 6D-E), which occurs in response to activation of RNase L (Burke et al., 2020).

We also observed that DHX9 knockdown led to increased PKR activation (Figure 6F-H). In this experiment, we used an A549 RNase L KO cell line co-expressing mApple-PKR and GFP-G3BP1, which allowed for visualization of stress granules following PKR mediated translation arrest (Corbet et al., 2022). The RNase L KO background cell line was used for these experiments because RNase L activation triggers the formation of G3BP1-enriched assemblies known as RLBs as opposed to stress granules (Burke et al., 2020), and can also lead to the phosphorylation of eIF2*α* (Burke et al., 2019) through the ribotoxic stress response (Karasik et al., 2026 RNA). We also measured PKR activation by immunostaining for p-PKR^Thr446^, a marker for activated PKR (Dey et al., 2014). We observed that PKR activation was enhanced in DHX9-depleted cells, as assessed by both the fluorescence intensity of p-PKR staining (Figure 6F-G) and the formation of stress granules (Figure 6F,6H). This observation is consistent with prior findings that DHX9 depletion can increase PKR activation (Zhou et al., 2024) suggesting that DHX9-mediated suppression of PKR activation may occur through limiting the formation of dRIFs.

Taken together, these results are consistent with prior work that DHX9 can limit dsRNA response signaling (Cottrell et al., 2024; Zhou et al., 2024). Since we observed that DHX9 can limit dRIF formation, and RNase L and PKR activation are positively correlated with dRIF formation, we suggest that a mechanism by which DHX9 may suppress RNase L and PKR activation is by limiting dRIF formation. However, it is clear that DHX9 can also limit dsRNA responses independently of dRIFs because DHX9 also limits IFN-I activation by RIG-I/MDA5, which are not enriched in these assemblies. We did not observe notable impacts on dsRNA response signaling from depletion of PACT or STAU1 (Figure S6A-G). ILF3 depletion resulted in decreased stress granule formation; however this was not associated with significantly decreased PKR activation (Figure S6E-F) and may be due to a previously reported role for ILF3 in regulating stress granule formation (Shiina & Nakayama, 2014).

### ZNF346 Enhances PKR Activation

Our analyses of dRIF components and dsRNA response signaling provided several observations that ZNF346 contributes to PKR activation. First, we observed a decrease in p-PKR staining in ZNF346-depleted cells (Figure 7A-B). Second, we observed a decrease in stress granule formation in ZNF346 depleted cells (Figure 7A,7C). Third, overexpression of an eGFP-ZNF346 protein resulted in co-localization of cytoplasmic eGFP-ZNF346 and p-PKR in cytoplasmic puncta and increased PKR activation in response to exogenous dsRNA (Figure 7F-G), but not in cells that were not treated with dsRNA (Figure 7F, S7B). Further, eGFP-ZNF346 overexpression enhanced translation repression compared to an eGFP vector, as assessed by a ribopuromycinylation assay (Figure 7H-I). These results suggest that ZNF346 enhances the activation of PKR. Moreover, ZNF346 knockdown did not appear to significantly change total PKR levels in cells (Figure S7A), indicating that ZNF346 enhances PKR activation by a mechanism other than by altering PKR levels.

**Figure 7.**
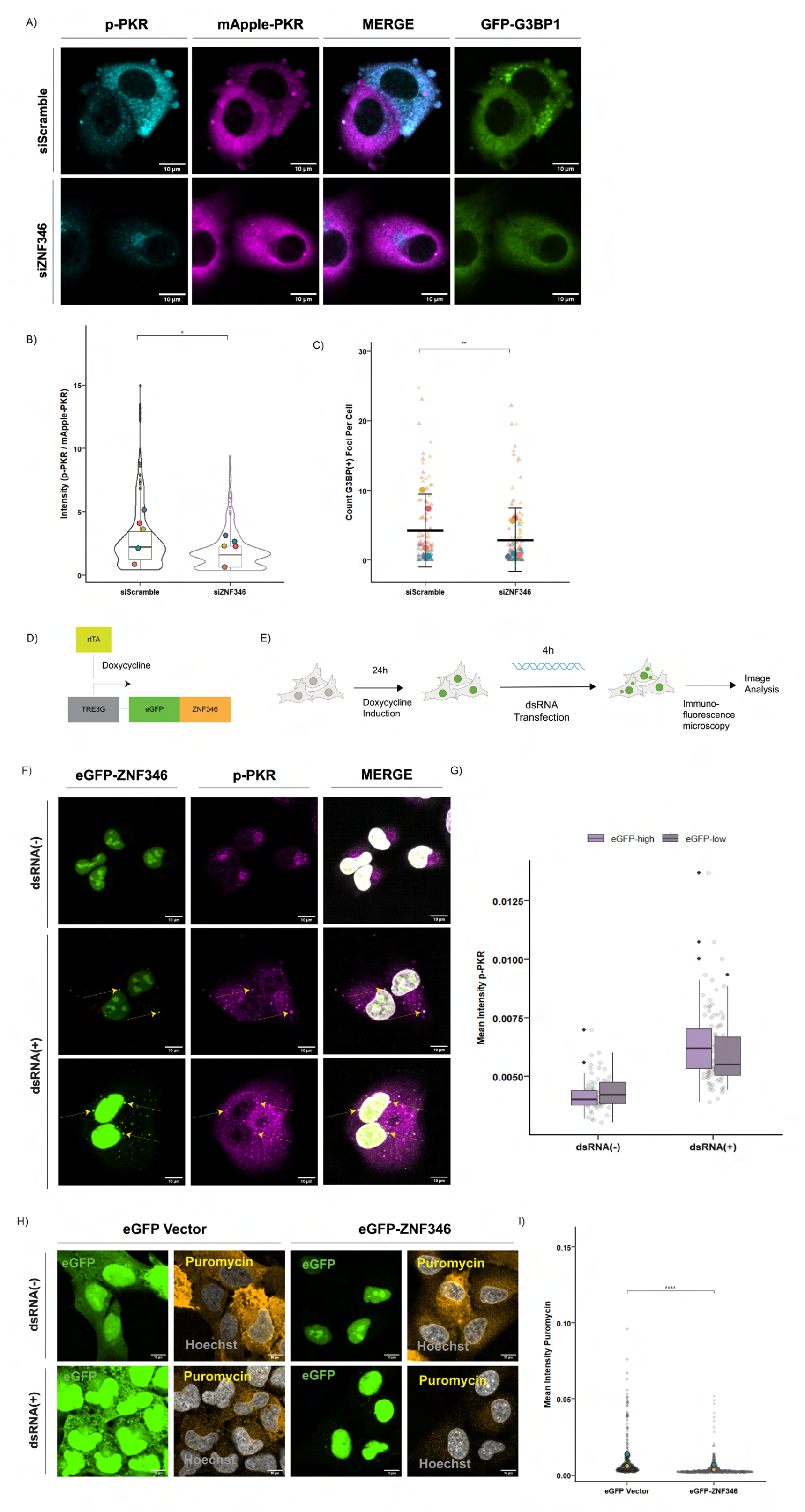
(**A**) Representative fluorescence microscopy images showing the relationship between PKR activation (visualized with an -p-PKR antibody), dRIF formation (denoted by mApple-PKR foci) and the presence of stress granules (denoted by GFP-G3BP1 foci) in dsRNA-transfected RNase L KO cells treated with siRNA targeting ZNF346 (siZNF346) or a non-targeting control (siScramble). (**B**) Quantification showing the mean intensity of p-PKR staining (normalized to mApple-PKR signal) in dsRNA-transfected RNase L KO cells treated with siRNA targeting ZNF346 (siZNF346) or a non-targeting control (siScramble). Statistical comparisons were made by performing an unpaired two-tailed t-test on per-image means. Data was obtained from 5 independent experiments and 300+ cells and 2+ fields-of-view per biological replicate and condition. (**C**) Quantification showing the percentage of stress granule-positive cells (proxied by GFP-G3BP1-positive foci) in dsRNA-transfected RNase L KO cells treated with siRNA targeting ZNF346 (siZNF346) or a non-targeting control (siScramble). Statistical comparisons were made by performing an unpaired two-tailed t-test on per-image means. Data was obtained from 5 independent experiments and 300+ cells and 5+ fields-of-view per biological replicate and condition. (**D**) Schematic for Tet-ON expression system for inducible eGFP-ZNF346 expression in A549 WT cells. (**E**) Temporal schematic for doxycycline-induced expression of eGFP-ZNF346 in A549 WT cells. (**F**) Representative fluorescence microscopy images showing doxycycline-induced expression of eGFP-ZNF346 and immunofluorescence staining for p-PKR in A549 WT cells treated with 2 ug/mL doxycycline with or without 400 ng/mL dsRNA treatment. (**G**) Quantification of the mean p-PKR immunofluorescence intensity in cells expressing eGFP-ZNF346 and treated with 2 ug/mL doxycycline with or without 400 ng/mL dsRNA treatment. Cells were binned according to eGFP nuclear fluorescence intensities, where eGFP-high cells are the top 50% and eGFP-low cells are the bottom 50% of single cells expressing eGFP-ZNF346. Circles represent individual data points. Data was obtained from 50+ cells for the no-dsRNA condition and 90+ cells for the dsRNA-treated condition. (**H**) Representative fluorescence microscopy images comparing the effects of expressing either eGFP tagged with V5 or eGFP-ZNF346 on translation arrest (proxied by *α*-puromycin staining intensity) in A549 WT cells transfected with 400 ng/mL dsRNA. (**I**) Quantification of mean -puromycin intensity in A549 WT cells expressing either eGFP tagged with V5 or eGFP-ZNF346 and transfected with 400 ng/mL dsRNA. Statistical comparisons were made using a Wilcoxon rank-sum test on individual data points. Data was obtained from 2 independent experiments and 150+ cells per biological replicate. All data in this figure shows cells transfected with 400 ng/mL dsRNA. All statistical analyses in this figure use the following p-value cutoffs: ^****^P ≤ 0.0001, ^***^P ≤ 0.001, ^**^P ≤ 0.01, ^*^P ≤ 0.05, n.s. P *>* 0.05.

**Figure 8.**
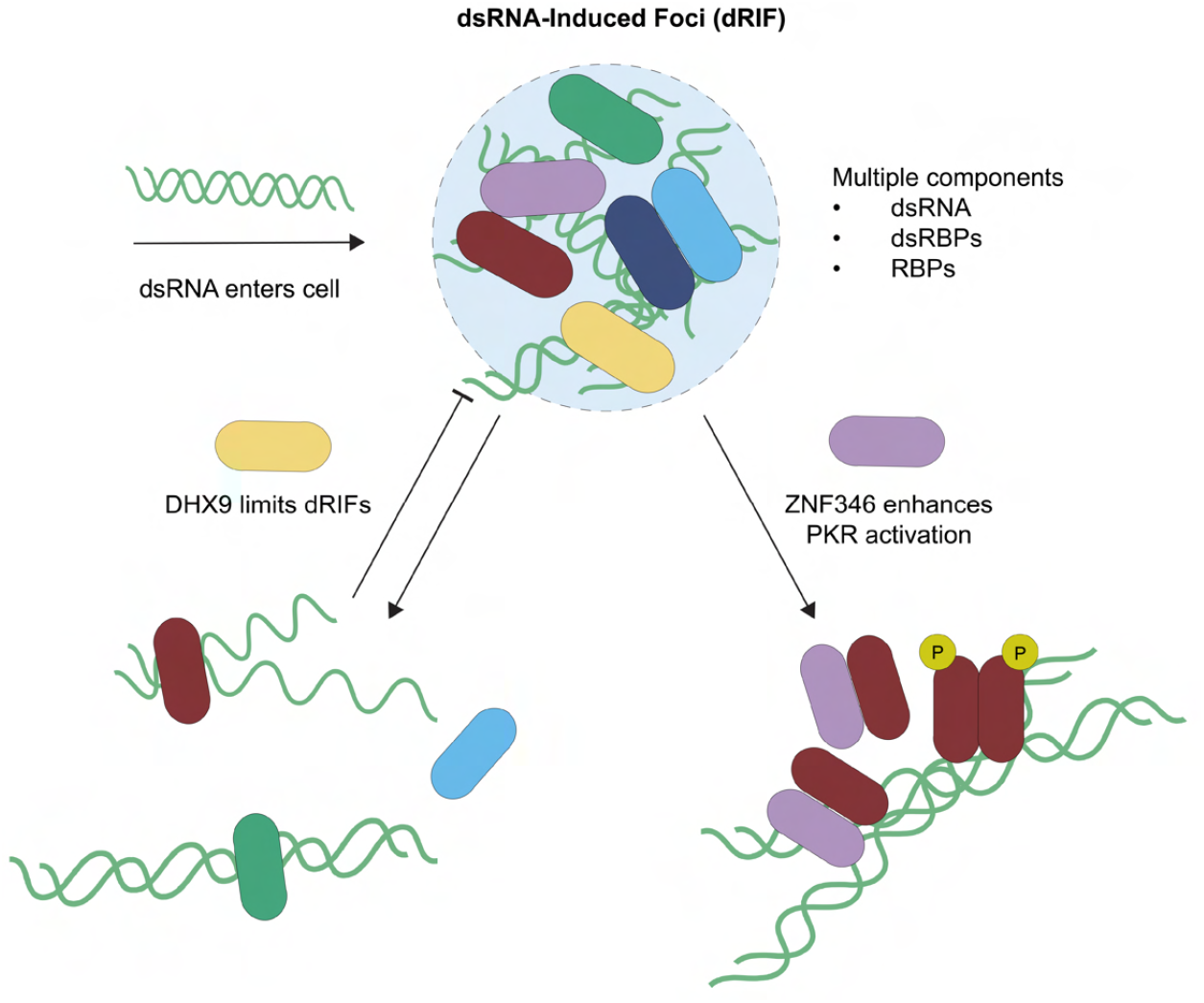
Graphical Abstract.

## DISCUSSION

In this work, we performed proteomics on dRIFs assembled from cell lysates, and validated both previously and newly annotated protein constituents of dRIFs. Our analysis suggests that these assemblies are more complex than previously described. Our confidence that these proteins are *bona fide* dRIF components is strengthened by our subsequent validation that the majority of these putative components localize to dRIFs *in situ*, or by expression of fluorescently tagged derivatives in cells. Specifically, we validated that cellular dRIFs can enrich ILF2, ILF3, NPM1, EEF1A, p62, ZNF346, and DDX18 (Figure 3 and 4), adding to previously identified components to provide an initial proteomic description of dRIFs (Figure S4C). Some of the new dRIF components are not known to bind dsRNA and are likely to be recruited to dRIFs by protein-protein interactions. For example, although ILF3 can bind dsRNA, ILF2 is not known to bind dsRNA and is presumably recruited to dRIFs by binding ILF3 (Bonczek et al., 2022). Since some of the newly annotated dRIF constituents are primary nuclear or nucleolar under homeostatic cellular conditions, this work underscores the importance of subcellular compartmentation of dsRNA in determining protein localization. For instance, the dsRBDs of ZNF346 are required for its nucleolar localization (M. Yang et al., 1999), but herein we demonstrate that increased cytoplasmic loads of dsRNA can promote its translocation to the cytosol.

Our results suggest that dRIF formation is correlated with activation of PKR and RNase L. This is suggested by the observations that DHX9 depletion increases dRIF formation and enhances PKR and RNase L signaling (Figure 5&6). These results are consistent with earlier work arguing DHX9 can limit cellular responses to dsRNA (Zhou et al., 2024) and that dRIF formation can enhance PKR or RNase L activation (Corbet et al., 2022; Cusic and Burke, 2024). However, it is also reasonable that DHX9 can modulate dsRNA signaling independently of dRIFs since DHX9 can unwind dsRNA (Lee & Pelletier, 2016). Intriguingly, the increase in MDA5/RIG-I activation was relatively weak relative to enhancement of RNase L and PKR activation in DHX9-depleted cells, suggesting that dRIFs primarily impact the OAS3-RNase L and PKR-eIF2*α* pathways, whereas regulation of MAVS signaling is largely independent of dRIFs. This is consistent with the absence of MDA5 from dRIFs, as reported both here (Figure 2) and previously (Corbet et al., 2022). This argues that dRIFs affect some but not all the cellular responses to dsRNA.

We have identified a granule related to dRIFs that forms during mitosis and persists into early anaphase, referred to as a mitotic dRIF. Our observations are consistent with reports of mitotic PKR foci forming elsewhere (Zappa et al., 2022), and with findings that PKR is activated by dsRNA-forming Alu elements during mitosis (Kim et al., 2014). Therefore, a parsimonious explanation for mitotic dRIF formation is that the release of IR-Alu repeat-containing RNAs during nuclear envelope breakdown in mitosis drives the assembly of mitotic dRIFs. These assemblies share many components with cytoplasmic dRIFs but differ in some intriguing manners. For example, we observe that DHX9 and ZNF346 are prevalent components of canonical dRIFs but are generally missing from mitotic dRIFs. In contrast, NPM1, which is seen in 19% of dRIFs induced by exogenous dsRNA (Figure 3), is present in essentially all mitotic dRIFs (Figure 4).

Since mitotic dRIFs have a different composition from canonical dRIFs, it remains to be determined if/how the formation of mitotic dRIFs influences the dsRNA response during this phase of the cell cycle. This is of interest since PKR has been suggested to be activated during mitosis and play a role in mitotic progression (Kim et al., 2014). However, the mechanisms by which cells prevent self-immunogenicity due to endogenous dsRNAs during mitosis require further investigation. We propose that mitotic dRIFs could serve to dampen innate immune responses during mitosis by sequestering immunogenic dsRNAs. One possibility is that mitotic dRIFs can be hijacked by viruses to evade immune recognition and infect daughter cells. Further, it will be interesting to determine whether these assemblies regulate aspects of cell cycle regulation, and how they may relate to adaptations in cancer cells that alter the levels of endogenous dsRNA and/or dsRBPs (Young et al., 2025).

In addition to differences between mitotic dRIFs and cytoplasmic dRIFs, several observations raise the possibility that there may be multiple subtypes of dRIFs that form in the cytoplasm with different effects on the dsRNA response. The possibility of subtypes of dRIFs is suggested by the observation here, and in prior work (Corbett et al., 2022), that some dRIF components are observed in only a fraction of these assemblies. For example, we observe that ILF3 is only present in 35% of dRIFs formed by exogenous dsRNA (Figure 3). Further analysis will inform whether altered dRIF composition contributes to the cell-to-cell heterogeneity observed in the dsRNA response.

We provide observations suggesting a role for ZNF346 in the cytosol as an activator of PKR. First, when we depleted ZNF346 from cells, we observed suppressed PKR activation following dsRNA transfection, as assessed by both reduced p-PKR levels and stress granule formation (Figure 7). Second, over-expression of ZNF346 leads to increased PKR activation and reduced puromycin incorporation after transfection of dsRNA (Figure 7). Since ZNF346 binds dsRNA (Burge et al., 2014), and co-IPs with PKR (Buljan et al., 2020; Cho et al., 2022; S. Li et al., 2011; Wang et al., 2023), a parsimonious model is that ZNF346 bound to dsRNA in the vicinity of PKR can also bind PKR to promote PKR auto-phosphorylation and activation. Interestingly, since ZNF346 is predominantly localized to the nucleolus, and has been implicated in apoptotic regulation through transcriptional control (Mallick & D’Mello, 2014), a role for ZNF346 in activating PKR would be a new function for this protein. ZNF346-mediated regulation of PKR would be expected to impact neuron homeostasis, as ZNF346 is abundantly expressed in the brain (T. Wu et al., 2018) and PKR dysregulation is linked to neurodegenerative disease (Marchal et al., 2014; Zu et al., 2020).

ZNF346 joins an emerging set of proteins that modulate the dsRNA response in the absence of any known catalytic activity, which includes PACT, ILF3, ZCCH3C, and OASL (Ahmad et al., 2025; Haque et al., 2023; M. Shi et al., 2025; Zhu et al., 2015). The high partitioning of ZNF346 into dRIFs relative to the free cytosol (Figure S7C) supports a model where localization of PKR regulators to dRIFs can modulate PKR activity. This highlights the importance of understanding what molecular complexes can form between these proteins on dsRNA and how that leads to activation of the critical dsRNA sensors, which will be an important area of future research.

## METHODS

### *In vitro* reconstitution of dRIFs (dRIF_IV_)

A549 PKR/RL dKO cells expressing mApple-mApple-PKR^K296R^ were cultured in 15 cm tissue culture plates to 80% confluency. 5 × 15cm plates were harvested per sample. On the day of harvest, cells were washed 2X with 5 mL PBS and detached from plates using a Corning cell lifter. Cells were transferred to a 15 mL falcon tube and centrifuged at 1300 RPM for 5 minutes to pellet the cells. PBS from the second wash was aspirated, followed by flash freezing of cells in liquid N2 and storage at -80C prior to lysis. Cells were thawed on ice for 5 min and lysed with a 25G needle in buffer composed of 50 mM Tris HCl pH 7.4, 100 mM KAc, 2 mM MgAc, 10 mM beta-mercapthoethanol, 1:5000 Antifoam B, and 0.65% n-Dodecyl-Beta-Maltoside detergent with one mini-tablet cOmplete Protease Inhibitor Cocktail per mL buffer. All subsequent centrifugation steps were performed at 4C. Cells were centrifuged at 1000xg for 5 minutes to remove cell debris. The supernatant was transferred and centrifuged at 10,000xg 1X for 10 minutes and 1X for 5 minutes to pellet nuclei. Poly(I:C) was added to lysed cells at 5 ug/mL and 10 ug/mL in pilot experiments, and incubated for 3 hours at 4C, to determine optimal concentrations for dRIF formation. For subsequent experiments shown in western blots and MS analysis, 6 ug/mL poly(I:C) and 3 hour incubation periods at 4C were used.

### Affinity purification of dRIF_IV_

Protein A Dynabeads (10008D Invitrogen) were pretreated in PBS with 1% DEPC for 1 hour at room temperature, then washed in PBS + 0.05% NP-50 for 5 minutes and washed 3X with lysis buffer at 4C. After poly(I:C) incubation, cell supernatants from the *in vitro* reconstitution steps were pre-cleared with Dynabeads at 4C for 30 minutes. 20 uL PKR antibody (Cell Signaling Technology 12297) was added to each sample followed by overnight incubation at 4C. dRIFs were then pelleted by centrifugation at 18,000xg 2X and resuspended in lysis buffer. Supernatants from the second spin were centrifuged 1X at 850xg for 5 minutes to pellet residual debris, and transferred to new microcentrifuge tubes. Supernatants were incubated for 3 hr at 4C with 60 uL Dynabeads per sample. Samples were washed with lysis buffer 3X and then 1X each with lysis buffer + 2 M urea, then 1X with lysis buffer + 300 mM KAc. Granules were eluted from Dynabeads in 20 uL SDS buffer with 15 mM betamercapthoethanol and 1X Protease/Phosphatase Inhibitor Cocktail (Cell Signaling Technology 5872). 2 uL of each sample was loaded onto a 4-12% Bis-Tris SDS-PAGE gel for western blot analysis.

### Sample preparation and mass spectrometry analysis

dRIF_IV_ samples were solubilized in 5% (w/v) SDS, 50 mM Tris-HCl, pH 8.5, 10 mM tris(2-carboxyethylphosphine) (TCEP) and 40 mM chloroacetamide by boiling at 95°C for 10 minutes. Each sample was digested using the SP3 method (Hughes et al., 2014).Two hundred micrograms carboxylate-functionalized speedbeads (GE Life Sciences) were added to the lysates. Addition of acetonitrile to 80% (v/v) caused the proteins to bind to precipitate on the beads. The beads were washed twice with 80% (v/v) ethanol and twice with 100% acetonitrile. Proteins were digested with Lys-C/Trypsin (Promega) overnight rotating at 37°C. Speedbeads were collected by centrifugation and then placed on a magnet to remove the digested peptides. The peptides were then desalted using an Oasis HLB cartridge (Waters) according to the manufacturer’s instructions and dried in a speedvac. Samples were suspended in 3% (v/v) acetonitrile/0.1% (v/v) trifluoroacetic acid and directly injected onto a reversed-phase C18 1.7 µm, 130 Å, 75 µm X 250 mm M-class column (Waters), using a Thermo Ultimate 3000 RSLCnano UPLC. Peptides were eluted at 300 nL/minute using a gradient from 2% to 15% acetonitrile over 40 min, then 15% to 40% in 5 minutes into a Q-Exactive HF-X mass spectrometer (Thermo Scientific). Precursor mass spectra (MS1) were acquired at a resolution of 120,000 from 350 to 1550 m/z with an automatic gain control (AGC) target of 3E6 and a maximum injection time of 50 milliseconds. Precursor peptide ion isolation width for MS2 fragment scans was 1.4 m/z, and the top 12 most intense ions were sequenced. All MS2 spectra were acquired at a resolution of 15,000 with higher energy collision dissociation (HCD) at 27% normalized collision energy. An AGC target of 1E5 and 100 milliseconds maximum injection time was used. Rawfiles were searched against the Uniprot Human database UP000005640 using Maxquant version 2.0.3.0 with cysteine carbamidomethylation as a fixed modification. Methionine oxidation and protein N-terminal acetylation were searched as variable modifications. All peptides and proteins were thresholded at a 1% false discovery rate (FDR).

### Cellular growth conditions

All cell lines were maintained in DMEM supplemented with 10% FBS and 1 % penicillin/streptomycin at 37°C and 5% CO2.

### Plasmids

pLV-EF1-mApple-PKR K296R and pLV-EF1-mApple-PKR plasmids were generated as previously described (Corbet et al., 2022). The eGFP-ZNF346 sequence was codon-optimized and synthesized by Twist Bioscience. For validation experiments using transiently transfected constructs, eGFP-ZNF346 was cloned into a pTwist CMV Puro vector from Twist Bioscience using the following primers: TTCCGAGCTCTC-GAATTCGCCACCATG (forward), GAGGCCTGCGGATCCTTAATCTTCCAGAGT (reverse). To generate cell lines stably expressing eGFP-ZNF346, the eGFP-ZNF346 fragment was cloned into either a pBS piggybac EF1-eGFP-BLAST (a gift from Carolyn Decker/Edward Courvan) or a PB-TRE-EGFP-EF1a-rtTA (Addgene 104454) backbone vector. eGFP from PB-TRE-EGFP-EF1a-rtTA was removed via enzyme restriction with NheI and XhoI prior to eGFP-ZNF346 co-ligation. eGFP from pBS piggybac EF1-eGFP-BLAST was removed via enzyme restriction with KpnI and XbaI prior to eGFP-ZNF346 co-ligation The eGFP-ZNF346 fragment was cloned into each vector using NEBuilder® HiFi DNA Assembly (NEB E5520). The following primers were used: GAAACTCGAGGGTACCGCCACCATG (forward pBS EF1-eGFP-ZNF346-BLAST), AAGCTGGGTCTAGATTAATCTTCCAGAG (reverse pBS EF1-eGFP-ZNF346-BLAST), TAAAGGTCTAGAGCTAGCGCCACCATG (forward PB-TRE-EGFP-ZNF346-EF1a-rtTA), AGGGATAGGCTTACCTTAATCTTCCAGAGT (reverse PB-TRE-EGFP-ZNF346-EF1a-rtTA), The K13-EF1a supperpiggybac transposase was originally a gift from the Michaal Ward lab.

### Cell line generation

A549 PKR/RL dKO mApple-PKR^K296R^, A549 PKR KO mApple-PKR^K296R^ GFP-G3BP1, and A549 PKR KO mApple-PKR GFP-G3BP1 cell lines were generated previously via lentiviral transduction against PKR and/or RL KO backgrounds (Corbet et al. 2023). The parental A549 cell line was provided by Dr. Christopher Sullivan (Burke et al., 2016). The A549 PKR KO, RNase L KO, and PKR/RNase L dKO cell lines were generated previously (Burke et al., 2019, 2020). Cell lines expressing EF1-eGFP-ZNF346 and Tet-On-3G-eGFP-ZNF346 were generated from an A549 WT parental cell line purchased from ATCC (CCL185). The eGFP-ZNF346 vectors were delivered to cells using a PiggyBac transposase system. Cells were co-transfected with Lipofectamine 3000, the donor plasmid and a K13 EF1 transposase vector. 2-3 days post-transfection, cells expressing the gene of interest were selected for by incubation with 2 ug/mL puromycin (Tet-On-3G-eGFP-ZNF346), blasticidin (EF1-eGFP-ZNF346) or by sorting for GFP(+) cells by flow cytometry after inducing fluorescent protein expression overnight using 1 ug/mL doxycyline. Antibiotic-containing media was replaced with standard growth media 2-3 days after antibiotic selection.

### dsRNA synthesis and labeling

Sequence-specific dsRNA was *in vitro*-transcribed from a plasmid with convergent T7 promoters (L4440-Y75B7AL.4), which contains a 0.9 kb insert sequence from C. elegans. L4440-Y75B7AL.4 was a gift from Hannah Fares at the University of Arizona. The T7 promoters and insert were PCR-amplified using the primer sequences TAATACGACTCACTATAGGGAGACC (forward) and TAATACGACTCACTATAGGGCGAATT (reverse) and PrimeSTAR HS DNA Polymerase pre-mix (Takara Bio R040A). PCR products were purified on a 1% agarose TAE gel using Qiagen gel extraction and cleanup reagents. *in vitro* transcription of the PCR amplicons was performed by incubation for 4 hours at 37C using a T7 Megascript kit (Ambion AM1334). Template DNA was degraded from the PCR reaction by incubation with Megascript Turbo DNase mix for 15 min at 37C. Complementary RNA strands were annealed by incubation at 75C for 5 minutes followed by incubation at RT for 10 minutes. Residual ssRNA was removed from the reaction by digestion with RNase I (Ambion) and RNase I buffer (100mM Tris Cl pH7.5, 50mM EDTA, 1.5M NaCl) for 1 hour at 37C. Reactions were purified both before and after RNase I treatment using a Megaclear Transcription Clean-Up kit (Invitrogen AM1908). Removal of ssRNA was assessed by running 500 ng of dsRNA product on a 1.2% agarose gel.

### Nucleic acid transfections and drug treatments

For poly(I:C) and dsRNA transfections, lipofectamine 2000 was used at a ratio of 1:2 nucleic acid:volume lipofectamine according to manufacturer’s instructions. Ribopuromycinylation assays were performed by incubating cells with puromycin (P8833; Sigma-Aldrich) at a working concentration of 10 ug/mL for 5 minutes at 37C prior to fixation with 4% PFA in PBS.

### Immunoblotting

For harvesting, cells were washed once with 1X PBS and lysed with Pierce RIPA Buffer (89900; Thermo Fisher Scientific), 1 mM beta-mercapthoethanol, and 1X Protease/Phosphatase Inhibitor Cocktail (Cell Signaling Technology 5872). Cells were incubated on ice for 20 minutes with vortexing every 10 minutes, then flash-frozen in liquid nitrogen and stored at -20C to promote cell lysis. 4X SDS loading buffer (200 mM Tris Cl pH 6.8, 8% SDS, 0.4% Bromophenol Blue, 40% glycerol) was added to cell lysates at a final 1X concentration, then incubated at 95C for 10 minutes. Samples were loaded on a 4-12% Bis-Tris-NuPAGE gel and transferred onto either a PVDF or nitrocellulose membrane. Blocking was performed with 5% BSA in TBS-T for 1 hour at room temperature, followed by primary antibody incubation with 5% BSA in TBS-T for 1 hour at room temperature or overnight at 4C. Membranes were washed three times with TBS-T and then incubated in secondary antibody with 5% BSA in 1X TBS-T for 1 hour at room temperature. Membranes were washed an additional three times with 1X TBS-T prior to antibody detection. For antibody detection, membranes were incubated with Pierce ECL Western blotting substrate for 5 minutes at room temperature and visualized using an Invitrogen iBright Imaging System.

### siRNA-mediated knockdown and experiments

siRNA knockdown was performed by reverse transfection. Cells were treated with 20 nM siRNA using Lipofectamine RNAiMAX (13778-150; Invitrogen). Specific siRNAs and catalog numbers are listed in Table S2. 5 µl lipofectamine was added to 150 µl Opti-MEM media per reaction. 20 nM siRNA was added to another tube with 150 µl Opti-MEM medium. Lipofectamine and siRNA mixtures were combined, then vortexed and incubated for 20 min at RT. 300 µl siRNA-lipofectamine mix was added per well and 2 ml media containing 250,000-300,000 cells per siRNA knockdown condition were added to the lipofection mixture in each well, at a final volume of 2.3 mL per well.

Cells were seeded onto a glass-bottom 96-well plate at 3500-4000 cells/well 24 hours after siRNA transfection and 24 hours prior to dsRNA transfection. Remaining cells were re-plated at a 1:3 ratio into a 6-well plate for immunoblot analysis and harvested upon 24 hr of re-plating. 4 hours after dsRNA transfection, mock-treated and siRNA-treated cells were washed once with PBS and then fixed with 4% paraformaldehyde in PBS for 10 minutes, then washed twice with PBS. Cells were simultaneously permeabilized and blocked with 0.1% Triton X-100 (AC327371000; Thermo Fisher Scientific) in 3% BSA (126593; Millipore Sigma) and PBS for either 1 hour at RT or overnight at 4C. Samples were incubated with primary antibody at either 1:500 or 1:1000 overnight at 4C, and washed 3X with PBS. Samples were incubated with secondary antibody at 1:1000 in 0.1% Triton X-100 in 3% BSA and PBS for 1 hour at room temperature and washed 3Xwith PBS prior to Hoechst staining (1:10,000) or coverslip mounting using Prolong Glass Antifade mountant with NucBlue (P36981; Thermo Fisher Scientific).

### Imaging parameters

Imaging was performed at room temperature using a Nikon SoRa spinning disk confocal microscope with either a 40X NA 1.15 water immersion objective or a 60X NA 1.27 water immersion objective at 1X or 2.8X magnification and a Hamamatsu OCRA Fusion BT sCMOS camera. NIS Elements software was used to acquire images. For siRNA knockdown experiments performed in 96-well plates, a custom pipeline created with Nikon JOBS software was used to record images automatically at 2-6 sites per well. Images were collected manually in 96-well plates in siRNA knockdown conditions where cells were sparse in the automatically assigned fields of view (FOVs). For manual imaging, FOVs were selected based on the presence of cells identified by Hoechst staining of nuclei in the DAPI channel. Automatically-obtained images with low contrast and/or fewer than 5 cells per FOV were excluded from analysis.

### Image analysis and quantification

Image processing was conducted using Fiji, where the sum intensity projection (Figure 5D-F) or sum intensity projection (all other figures) was obtained from a series of z-slices from each image. Qualitative images shown were assigned minimum and maximum display values and converted to RGB color in ImageJ 2.14.0, Java 1.8.0_322. CellProfiler version 4.2.8 was used for image analysis (granule quantification, obtainment of fluorescence intensity values) of maximum intensity-projected images. The Cellpose plugin for CellProfiler was used for segmentation of the cytoplasm. Fluorescence intensity values from CellProfiler output files were used to calculate ratiometric measurements and partition coefficients. Nucleocytoplasmic intensity ratios were calculated by dividing the mean nuclear fluorescence intensity by the mean cytosolic fluorescence intensity.

### Statistical analysis

MS data: Analysis of “proteinGroups.txt” output files from MaxQuant was performed in RStudio using the DEP2 package. Data was pre-processed by filtering out reverse sequences and common contaminants including keratins. Raw intensity values for each protein were log2 transformed and normalized via median normalization with the proDA package. Values were imputed under a missing not at random (MNAR) assumption by a minimum probability function centered around a q-value of 0.01. Differentially enriched proteins were defined as proteins with a log2 fold change greater than or equal to 1 with respect to either a PKR KO or poly(I:C)-background control. False discovery rates were determined using a Benjamini-Hochberg procedure.

Fluorescence microscopy data: Statistical analysis was performed in RStudio 2023.06.0 using the stat_compare_means function. The data from each siRNA knockdown condition was tested for normality using a Q-Q plot and Shapiro-Wilk test prior to selection of a parametric or non-parametric statistical test. Fluorescence intensities for readouts of dsRNA response signaling were normalized to a no-dsRNA control treated with non-targeting siRNA.

## Supporting information

Supplemental Table (Reagents & Other Materials)

## ACKNOWLEDGMENTS

We thank the Proteomics and Mass Spectrometry Core Facility (RRID:SCR_ 018992) in the Department of Biochemistry at the University of Colorado Boulder for sample preparation, data acquisition and analysis using the Thermo Q-Exactive HF-X mass spectrometer, funded by NIH Grant S10-OD025267. We also thank Dr. Joseph Dragavon (BioFrontiers Institute Advanced Light Microscopy Core Facility), Theresa Nahreini, and Emily Proksch (Biochemistry Cell Culture Facility (RRID:SCR_018988)) in the Department of Biochemistry at the University of Colorado Boulder. The research from this manuscript is supported by Howard Hughes Medical Institute (HHMI) funds (R. Parker).

## REFERENCES

Ahmad, S., Zou, T., Hwang, J., Zhao, L., Wang, X., Davydenko, A., Buchumenski, I., Zhuang, P., Fishbein, A. R., Capcha-Rodriguez, D., Orgel, A., Levanon, E. Y., Myong, S., Chou, J., Meyerson, M., & Hur, S. (2025). PACT prevents aberrant activation of PKR by endogenous dsRNA without sequestration. Nature Communications, 16(1), 3325. 10.1038/s41467-025-58433-x

Berke, I. C., Yu, X., Modis, Y., & Egelman, E. H. (2012). MDA5 assembles into a polar helical filament on dsRNA. Proceedings of the National Academy of Sciences, 109(45), 18437–18441. 10.1073/pnas.1212186109

Bonczek, O., Wang, L., Gnanasundram, S. V., Chen, S., Haronikova, L., Zavadil-Kokas, F., & Vojtesek, B. (2022). DNA and RNA Binding Proteins: From Motifs to Roles in Cancer. International Journal of Molecular Sciences, 23(16), 9329. 10.3390/ijms23169329

Buljan, M., Ciuffa, R., Van Drogen, A., Vichalkovski, A., Mehnert, M., Rosenberger, G., Lee, S., Varjosalo, M., Pernas, L. E., Spegg, V., Snijder, B., Aebersold, R., & Gstaiger, M. (2020). Kinase Interaction Network Expands Functional and Disease Roles of Human Kinases. Molecular Cell, 79(3), 504-520.e9. 10.1016/j.molcel.2020.07.001

Burge, R. G., Martinez-Yamout, M. A., Dyson, H. J., & Wright, P. E. (2014). Structural characterization of interactions between the double-stranded RNA-binding zinc finger protein JAZ and nucleic acids. Biochemistry, 53(9), 1495–1510. 10.1021/bi401675h

Burke, J. M., Kincaid, R. P., Nottingham, R. M., Lambowitz, A. M., & Sullivan, C. S. (2016). DUSP11 activity on triphosphorylated transcripts promotes Argonaute association with noncanonical viral microRNAs and regulates steady-state levels of cellular noncoding RNAs. Genes & Development, 30(18), 2076–2092. 10.1101/gad.282616.116

Burke, J. M., Lester, E. T., Tauber, D., & Parker, R. (2020). RNase L promotes the formation of unique ribonucleoprotein granules distinct from stress granules. The Journal of Biological Chemistry, 295(6), 1426–1438. 10.1074/jbc.RA119.011638

Burke, J. M., Moon, S. L., Matheny, T., & Parker, R. (2019). RNase L Reprograms Translation by Widespread mRNA Turnover Escaped by Antiviral mRNAs. Molecular Cell, 75(6), 1203-1217.e5. 10.1016/j.molcel.2019.07.029

Chan, C.-P., & Jin, D.-Y. (2022). Cytoplasmic RNA sensors and their interplay with RNA-binding partners in innate antiviral response: Theme and variations. RNA (New York, N.Y.), 28(4), 449–477. 10.1261/rna.079016.121

Chan, W. Y., Liu, Q. R., Borjigin, J., Busch, H., Rennert, O. M., Tease, L. A., & Chan, P. K. (1989). Characterization of the cDNA encoding human nucleophosmin and studies of its role in normal and abnormal growth. Biochemistry, 28(3), 1033–1039. 10.1021/bi00429a017

Chen, T., Brownawell, A. M., & Macara, I. G. (2004). Nucleocytoplasmic Shuttling of JAZ, a New Cargo Protein for Exportin-5. Molecular and Cellular Biology, 24(15), 6608–6619. 10.1128/MCB.24.15.6608-6619.2004

Chen, Y. G., & Hur, S. (2022). Cellular origins of dsRNA, their recognition and consequences. Nature Reviews. Molecular Cell Biology, 23(4), 286–301. 10.1038/s41580-021-00430-1

Cho, N. H., Cheveralls, K. C., Brunner, A.-D., Kim, K., Michaelis, A. C., Raghavan, P., Kobayashi, H., Savy, L., Li, J. Y., Canaj, H., Kim, J. Y. S., Stewart, E. M., Gnann, C., McCarthy, F., Cabrera, J. P., Brunetti, R. M., Chhun, B. B., Dingle, G., Hein, M. Y., … Leonetti, M. D. (2022). OpenCell: Endogenous tagging for the cartography of human cellular organization. Science, 375(6585), eabi6983. 10.1126/science.abi6983

Chukwurah, E., Farabaugh, K. T., Guan, B., Ramakrishnan, P., & Hatzoglou, M. (2021). A tale of two proteins: PACT and PKR and their roles in inflammation. The FEBS Journal, 288(22), 6365–6391. 10.1111/febs.15691

Corbet, G. A., Burke, J. M., Bublitz, G. R., Tay, J. W., & Parker, R. (2022). dsRNA-induced condensation of antiviral proteins modulates PKR activity. Proceedings of the National Academy of Sciences of the United States of America, 119(33), e2204235119. 10.1073/pnas.2204235119

Cottrell, K. A., Ryu, S., Pierce, J. R., Soto Torres, L., Bohlin, H. E., Schab, A. M., & Weber, J. D. (2024). Induction of Viral Mimicry Upon Loss of DHX9 and ADAR1 in Breast Cancer Cells. Cancer Research Communications, 4(4), 986–1003. 10.1158/2767-9764.CRC-23-0488

Cusic, R., & Burke, J. M. (2024). Condensation of RNase L promotes its rapid activation in response to viral infection in mammalian cells. Science Signaling, 17(837), eadi9844. 10.1126/scisignal.adi9844

Dey, M., Mann, B. R., Anshu, A., & Mannan, M. A. (2014). Activation of Protein Kinase PKR Requires Dimerization-induced cis-Phosphorylation within the Activation Loop. Journal of Biological Chemistry, 289(9), 5747–5757. 10.1074/jbc.M113.527796

Dorrity, T. J., Shin, H., Wiegand, K. A., Aruda, J., Closser, M., Jung, E., Gertie, J. A., Leone, A., Polfer, R., Culbertson, B., Yu, L., Wu, C., Ito, T., Huang, Y., Steckelberg, A. L., Wichterle, H., & Chung, H. (2023). Long 3’UTRs predispose neurons to inflammation by promoting immunostimulatory double-stranded RNA formation. Science immunology, 8(88), eadg2979. 10.1126/sciimmunol.adg2979

Freibaum, B. D., Messing, J., Yang, P., Kim, H. J., & Taylor, J. P. (2021). High-fidelity reconstitution of stress granules and nucleoli in mammalian cellular lysate. The Journal of cell biology, 220(3), e202009079. 10.1083/jcb.202009079

Gal-Ben-Ari, S., Barrera, I., Ehrlich, M., & Rosenblum, K. (2018). PKR: A Kinase to Remember. Frontiers in Molecular Neuroscience, 11, 480. 10.3389/fnmol.2018.00480

Gantier, M. P., & Williams, B. R. (2007). The response of mammalian cells to double-stranded RNA. Cytokine & growth factor reviews, 18(5-6), 363–371. 10.1016/j.cytogfr.2007.06.016

Haque, N., Will, A., Cook, A. G., & Hogg, J. R. (2023). A network of DZF proteins controls alternative splicing regulation and fidelity. Nucleic Acids Research, 51(12), 6411–6429. 10.1093/nar/gkad351

Harashima, A., Guettouche, T., & Barber, G. N. (2010). Phosphorylation of the NFAR proteins by the dsRNA-dependent protein kinase PKR constitutes a novel mechanism of translational regulation and cellular defense. Genes & Development, 24(23), 2640–2653. 10.1101/gad.1965010

Hartmann, G. (2017). Nucleic Acid Immunity. Advances in Immunology, 133, 121–169. 10.1016/bs.ai.2016.11.001

Hughes, C. S., Foehr, S., Garfield, D. A., Furlong, E. E., Steinmetz, L. M., & Krijgsveld, J. (2014). Ultrasensitive proteome analysis using paramagnetic bead technology. Molecular Systems Biology, 10(10), 757. 10.15252/msb.20145625

Jeon, J., & Kim, Y. (2025). Toward unraveling molecular grammars for dsRNA-binding proteins: Substrate recognition to binding mechanisms. BMB Reports, 58(11), 451–466. 10.5483/BMBRep.2025-0067

Kedersha, N., Panas, M. D., Achorn, C. A., Lyons, S., Tisdale, S., Hickman, T., Thomas, M., Lieberman, J., McInerney, G. M., Ivanov, P., & Anderson, P. (2016). G3BP-Caprin1-USP10 complexes mediate stress granule condensation and associate with 40S subunits. The Journal of Cell Biology, 212(7), 845–860. 10.1083/jcb.201508028

Keeble, A. H., Khan, Z., Forster, A., & James, L. C. (2008). TRIM21 is an IgG receptor that is structurally, thermodynamically, and kinetically conserved. Proceedings of the National Academy of Sciences, 105(16), 6045–6040. 10.1073/pnas.0800159105

Kim, Y., Lee, J. H., Park, J.-E., Cho, J., Yi, H., & Kim, V. N. (2014). PKR is activated by cellular dsRNAs during mitosis and acts as a mitotic regulator. Genes & Development, 28(12), 1310–1322. 10.1101/gad.242644.114

Lee, T., & Pelletier, J. (2016). The biology of DHX9 and its potential as a therapeutic target. Oncotarget, 7(27), 42716–42739. 10.18632/oncotarget.8446

Lester, E., Van Alstyne, M., McCann, K. L., Reddy, S., Cheng, L. Y., Kuo, J., Pratt, J., & Parker, R. (2023). Cytosolic condensates rich in polyserine define subcellular sites of tau aggregation. Proceedings of the National Academy of Sciences, 120(3), e2217759120. 10.1073/pnas.2217759120

Li, S., Wang, L., Berman, M., Kong, Y.-Y., & Dorf, M. E. (2011). Mapping a Dynamic Innate Immunity Protein Interaction Network Regulating Type I Interferon Production. Immunity, 35(3), 426–440. 10.1016/j.immuni.2011.06.014

Li, Y., Banerjee, S., Wang, Y., Goldstein, S. A., Dong, B., Gaughan, C., Silverman, R. H., & Weiss, S. R. (2016). Activation of RNase L is dependent on OAS3 expression during infection with diverse human viruses. Proceedings of the National Academy of Sciences of the United States of America, 113(8), 2241–2246. 10.1073/pnas.1519657113

Liddicoat, B. J., Piskol, R., Chalk, A. M., Ramaswami, G., Higuchi, M., Hartner, J. C., Li, J. B., Seeburg, P. H., & Walkley, C. R. (2015). RNA editing by ADAR1 prevents MDA5 sensing of endogenous dsRNA as nonself. Science (New York, N.Y.), 349(6252), 1115–1120. 10.1126/science.aac7049

Lim, J., Lee, N., Ju, S., Kim, J., Mun, S., Jeon, M., Lee, Y., Lee, S.-H., Ku, J., Kim, S., Bae, S., Kim, J.-S., & Kim, Y. (2025). Cellular dsRNA interactome captured by K1 antibody reveals the regulatory map of exogenous RNA sensing. Communications Biology, 8(1), 389. 10.1038/s42003-025-07807-4

Liu, C.-X., Li, X., Nan, F., Jiang, S., Gao, X., Guo, S.-K., Xue, W., Cui, Y., Dong, K., Ding, H., Qu, B., Zhou, Z., Shen, N., Yang, L., & Chen, L.-L. (2019). Structure and Degradation of Circular RNAs Regulate PKR Activation in Innate Immunity. Cell, 177(4), 865-880.e21. 10.1016/j.cell.2019.03.046

Liu, S., Cai, X., Wu, J., Cong, Q., Chen, X., Li, T., Du, F., Ren, J., Wu, Y.-T., Grishin, N. V., & Chen, Z. J. (2015). Phosphorylation of innate immune adaptor proteins MAVS, STING, and TRIF induces IRF3 activation. Science, 347(6227), aaa2630. 10.1126/science.aaa2630

Maelfait, J., Liverpool, L., & Rehwinkel, J. (2020). Nucleic Acid Sensors and Programmed Cell Death. Journal of Molecular Biology, 432(2), 552–568. 10.1016/j.jmb.2019.11.016

Mallick, S., & D’Mello, S. R. (2014). JAZ (Znf346), a SIRT1-interacting protein, protects neurons by stimulating p21 (WAF/CIP1) protein expression. The Journal of Biological Chemistry, 289(51), 35409–35420. 10.1074/jbc.M114.597575

Marchal, J. A., Lopez, G. J., Peran, M., Comino, A., Delgado, J. R., García-García, J. A., Conde, V., Aranda, F. M., Rivas, C., Esteban, M., & Garcia, M. A. (2014). The impact of PKR activation: From neurodegeneration to cancer. FASEB Journal, 28(5), 1965–1974. 10.1096/fj.13-248294

Martel, C., Macchi, P., Furic, L., Kiebler, M. A., & Desgroseillers, L. (2006). Staufen1 is imported into the nucleolus via a bipartite nuclear localization signal and several modulatory determinants. Biochemical Journal, 393(1), 245–254. 10.1042/BJ20050694

Mellacheruvu, D., Wright, Z., Couzens, A. L., Lambert, J. P., St-Denis, N. A., Li, T., Miteva, Y. V., Hauri, S., Sardiu, M. E., Low, T. Y., Halim, V. A., Bagshaw, R. D., Hubner, N. C., Al-Hakim, A., Bouchard, A., Faubert, D., Fermin, D., Dunham, W. H., Goudreault, M., Lin, Z. Y., … Nesvizhskii, A. I. (2013). The CRAPome: a contaminant repository for affinity purification-mass spectrometry data. Nature methods, 10(8), 730–736. 10.1038/nmeth.2557

Milstead, R. A., Link, C. D., Xu, Z., & Hoeffer, C. A. (2023). TDP-43 knockdown in mouse model of ALS leads to dsRNA deposition, gliosis, and neurodegeneration in the spinal cord. Cerebral cortex, 33(10), 5808–5816. 10.1093/cercor/bhac461

Nabeel-Shah, S., Lee, H., Ahmed, N., Burke, G. L., Farhangmehr, S., Ashraf, K., Pu, S., Braunschweig, U., Zhong, G., Wei, H., Tang, H., Yang, J., Marcon, E., Blencowe, B. J., Zhang, Z., & Greenblatt, J. F. (2022). SARS-CoV-2 nucleocapsid protein binds host mRNAs and attenuates stress granules to impair host stress response. iScience, 25(1), 103562. 10.1016/j.isci.2021.103562

Nourreddine, S., Lavoie, G., Paradis, J., Ben El Kadhi, K., Méant, A., Aubert, L., Grondin, B., Gendron, P., Chabot, B., Bouvier, M., Carreno, S., & Roux, P. P. (2020). NF45 and NF90 Regulate Mitotic Gene Expression by Competing with Staufen-Mediated mRNA Decay. Cell Reports, 31(7), 107660. 10.1016/j.celrep.2020.107660

Pang, Q., Christianson, T. A., Koretsky, T., Carlson, H., David, L., Keeble, W., Faulkner, G. R., Speckhart, A., & Bagby, G. C. (2003). Nucleophosmin Interacts with and Inhibits the Catalytic Function of Eukaryotic Initiation Factor 2 Kinase PKR. Journal of Biological Chemistry, 278(43), 41709–41717. 10.1074/jbc.M301392200

Peisley, A., Lin, C., Wu, B., Orme-Johnson, M., Liu, M., Walz, T., & Hur, S. (2011). Cooperative assembly and dynamic disassembly of MDA5 filaments for viral dsRNA recognition. Proceedings of the National Academy of Sciences, 108(52), 21010–21015. 10.1073/pnas.1113651108

Pfaller, C. K., Donohue, R. C., Nersisyan, S., Brodsky, L., & Cattaneo, R. (2018). Extensive editing of cellular and viral double-stranded RNA structures accounts for innate immunity suppression and the proviral activity of ADAR1p150. PLoS Biology, 16(11), e2006577. 10.1371/journal.pbio.2006577

Prasanth, K. V., Sacco-Bubulya, P. A., Prasanth, S. G., & Spector, D. L. (2003). Sequential Entry of Components of Gene Expression Machinery into Daughter Nuclei. Molecular Biology of the Cell, 14(3), 1043–1057. 10.1091/mbc.e02-10-0669

Rai, A. K., Chen, J.-X., Selbach, M., & Pelkmans, L. (2018). Kinase-controlled phase transition of membraneless organelles in mitosis. Nature, 559(7713), 211–216. 10.1038/s41586-018-0279-8

Rodriguez, S., Sahin, A., Schrank, B. R., Al-Lawati, H., Costantino, I., Benz, E., Fard, D., Albers, A. D., Cao, L., Gomez, A. C., Evans, K., Ratti, E., Cudkowicz, M., Frosch, M. P., Talkowski, M., Sorger, P. K., Hyman, B. T., & Albers, M. W. (2021). Genome-encoded cytoplasmic double-stranded RNAs, found in C9ORF72 ALS-FTD brain, propagate neuronal loss. Science translational medicine, 13(601), eaaz4699. 10.1126/scitranslmed.aaz4699

Sanchez David, R. Y., Combredet, C., Najburg, V., Millot, G. A., Beauclair, G., Schwikowski, B., Léger, T., Camadro, J.-M., Jacob, Y., Bellalou, J., Jouvenet, N., Tangy, F., & Komarova, A. V. (2019). LGP2 binds to PACT to regulate RIG-I- and MDA5-mediated antiviral responses. Science Signaling, 12(601), eaar3993. 10.1126/scisignal.aar3993

Sharp, T. V., Xiao, Q., Justesen, J., Gewert, D. R., & Clemens, M. J. (1995). Regulation of the interferon-inducible protein kinase PKR and (2’-5’)oligo(adenylate) synthetase by a catalytically inactive PKR mutant through competition for double-stranded RNA binding. European journal of biochemistry, 230(1), 97–103. 10.1111/j.1432-1033.1995.0097i.x

Shi, M., Jiang, T., Zhang, M., Li, Q., Liu, K., Lin, N., Wang, X., Jiang, A., Gao, Y., Wang, Y., Liu, S., Zhang, L., Li, D., & Gao, P. (2025). Nucleic-acid-induced ZCCHC3 condensation promotes broad innate immune responses. Molecular Cell, 85(5), 962-975.e7. 10.1016/j.molcel.2025.01.027

Shi, X., Li, Y., Zhou, H., Hou, X., Yang, J., Malik, V., Faiola, F., Ding, J., Bao, X., Modic, M., Zhang, W., Chen, L., Mahmood, S. R., Apostolou, E., Yang, F.-C., Xu, M., Xie, W., Huang, X., Chen, Y., & Wang, J. (2024). DDX18 coordinates nucleolus phase separation and nuclear organization to control the pluripotency of human embryonic stem cells. Nature Communications, 15(1), 10803. 10.1038/s41467-024-55054-8

Shiina, N., & Nakayama, K. (2014). RNA granule assembly and disassembly modulated by nuclear factor associated with double-stranded RNA 2 and nuclear factor 45. The Journal of Biological Chemistry, 289(30), 21163–21180. 10.1074/jbc.M114.556365

Sinigaglia, K., Cherian, A., Du, Q., Lacovich, V., Vukić, D., Melicherová, J., Linhartova, P., Zerad, L., Stejskal, S., Malik, R., Prochazka, J., Bondurand, N., Sedláček, R., O’Connell, M. A., & Keegan, L. P. (2024). An ADAR1 dsRBD3-PKR kinase domain interaction on dsRNA inhibits PKR activation. Cell Reports, 43(8), 114618. 10.1016/j.celrep.2024.114618

Tauber, D., Tauber, G., Khong, A., Van Treeck, B., Pelletier, J., & Parker, R. (2020). Modulation of RNA Condensation by the DEAD-Box Protein eIF4A. Cell, 180(3), 411-426.e16. 10.1016/j.cell.2019.12.031

Viranaicken, W., Gasmi, L., Chaumet, A., Durieux, C., Georget, V., Denoulet, P., & Larcher, J.-C. (2011). L-Ilf3 and L-NF90 traffic to the nucleolus granular component: Alternatively-spliced exon 3 encodes a nucleolar localization motif. PloS One, 6(7), e22296. 10.1371/journal.pone.0022296

Wang, X.-W., Madeddu, L., Spirohn, K., Martini, L., Fazzone, A., Becchetti, L., Wytock, T. P., Kovács, I. A., Balogh, O. M., Benczik, B., Pétervári, M., Ágg, B., Ferdinandy, P., Vulliard, L., Menche, J., Colonnese, S., Petti, M., Scarano, G., Cuomo, F., … Liu, Y.-Y. (2023). Assessment of community efforts to advance network-based prediction of protein–protein interactions. Nature Communications, 14(1), 1582. 10.1038/s41467-023-37079-7

Wolkowicz, U. M., & Cook, A. G. (2012). NF45 dimerizes with NF90, Zfr and SPNR via a conserved domain that has a nucleotidyltransferase fold. Nucleic Acids Research, 40(18), 9356–9368. 10.1093/nar/gks696

Wu, B., Peisley, A., Richards, C., Yao, H., Zeng, X., Lin, C., Chu, F., Walz, T., & Hur, S. (2013). Structural basis for dsRNA recognition, filament formation, and antiviral signal activation by MDA5. Cell, 152(1–2), 276–289. 10.1016/j.cell.2012.11.048

Wu, J., & Chen, Z. J. (2014). Innate Immune Sensing and Signaling of Cytosolic Nucleic Acids. Annual Review of Immunology, 32(1), 461–488. 10.1146/annurev-immunol-032713-120156

Wu, T., Lin, Y., & Xie, Z. (2018). MicroRNA-1247 inhibits cell proliferation by directly targeting ZNF346 in childhood neuroblastoma. Biological Research, 51(1), 13. 10.1186/s40659-018-0162-y

Yang, M., May, W. S., & Ito, T. (1999). JAZ Requires the Double-stranded RNA-binding Zinc Finger Motifs for Nuclear Localization. Journal of Biological Chemistry, 274(39), 27399–27406. 10.1074/jbc.274.39.27399

Yang, P., Mathieu, C., Kolaitis, R.-M., Zhang, P., Messing, J., Yurtsever, U., Yang, Z., Wu, J., Li, Y., Pan, Q., Yu, J., Martin, E. W., Mittag, T., Kim, H. J., & Taylor, J. P. (2020). G3BP1 Is a Tunable Switch that Triggers Phase Separation to Assemble Stress Granules. Cell, 181(2), 325-345.e28. 10.1016/j.cell.2020.03.036

Young, A. A., Juhler, I. G., Pierce, J. R., Bohlin, H. E., Harper, H. A., Onishile, D. S., Chua, R. N., Liu, M. E., Gardner, E. N., Elzey, B. D., & Cottrell, K. A. (2025). PACT suppresses PKR activation through dsRNA binding and dimerization, and is a therapeutic target for triple-negative breast cancer. RNA, 31(11), 1599–1618. 10.1261/rna.080637.125

Zappa, F., Muniozguren, N. L., Wilson, M. Z., Costello, M. S., Ponce-Rojas, J. C., & Acosta-Alvear, D. (2022). Signaling by the integrated stress response kinase PKR is fine-tuned by dynamic clustering. The Journal of Cell Biology, 221(7), e202111100. 10.1083/jcb.202111100

Zhou, Y., Panhale, A., Shvedunova, M., Balan, M., Gomez-Auli, A., Holz, H., Seyfferth, J., Helmstädter, M., Kayser, S., Zhao, Y., Erdogdu, N. U., Grzadzielewska, I., Mittler, G., Manke, T., & Akhtar, A. (2024). RNA damage compartmentalization by DHX9 stress granules. Cell, 187(7), 1701-1718.e28. 10.1016/j.cell.2024.02.028

Zhu, J., Ghosh, A., & Sarkar, S. N. (2015). OASL-a new player in controlling antiviral innate immunity. Current Opinion in Virology, 12, 15–19. 10.1016/j.coviro.2015.01.010

Zu, T., Guo, S., Bardhi, O., Ryskamp, D. A., Li, J., Khoramian Tusi, S., Engelbrecht, A., Klippel, K., Chakrabarty, P., Nguyen, L., Golde, T. E., Sonenberg, N., & Ranum, L. P. W. (2020). Metformin inhibits RAN translation through PKR pathway and mitigates disease in C9orf72 ALS/FTD mice. Proceedings of the National Academy of Sciences of the United States of America, 117(31), 18591–18599. 10.1073/pnas.2005748117

