## Supplementary Figures for "Analysis of dRIF Composition Identifies ZNF346 as a Regulator of PKR Activation"

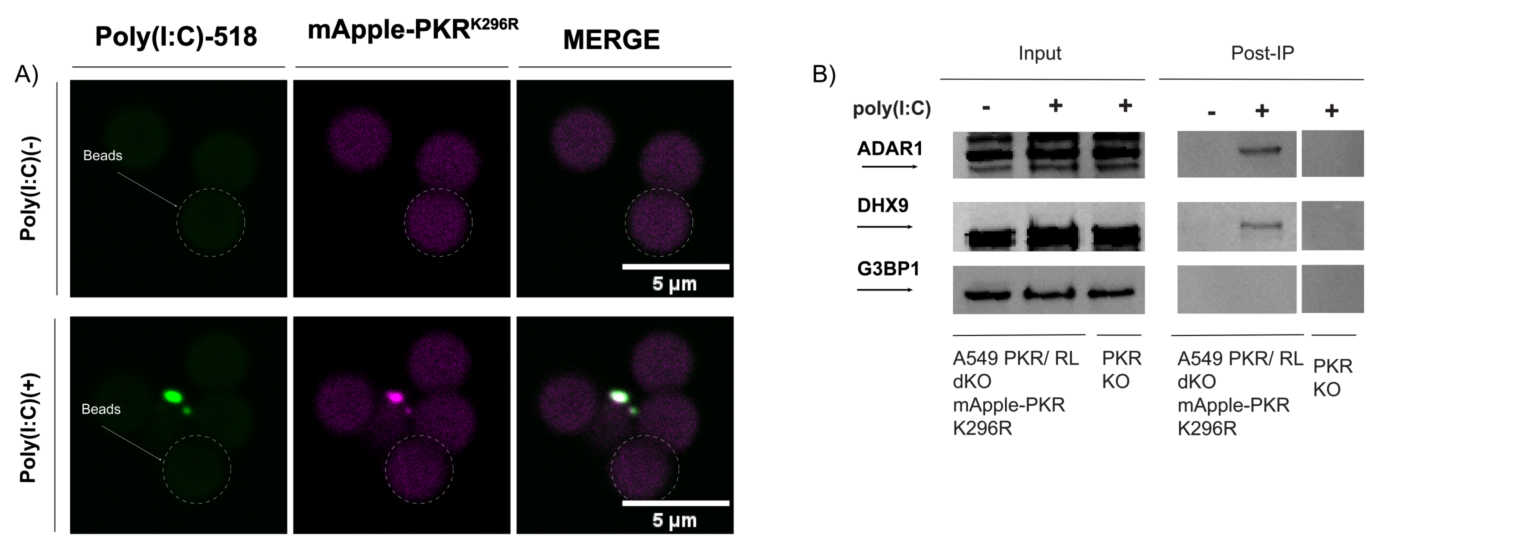


**Figure S1.** (**A**) Representative fluorescence microscopy images assessing the stability of $dRIF_{IV}$ through the terminal steps of the co-IP process (24+ hours). (**B**) Western blot analysis assessing the enrichment of known dRIF constituents (mApple-PKR, ADAR1, DHX9) and the exclusion of proteins known to form independent granules (G3BP1) in immunopurified $dRIF_{IV}$ compared to a PKR KO negative control.

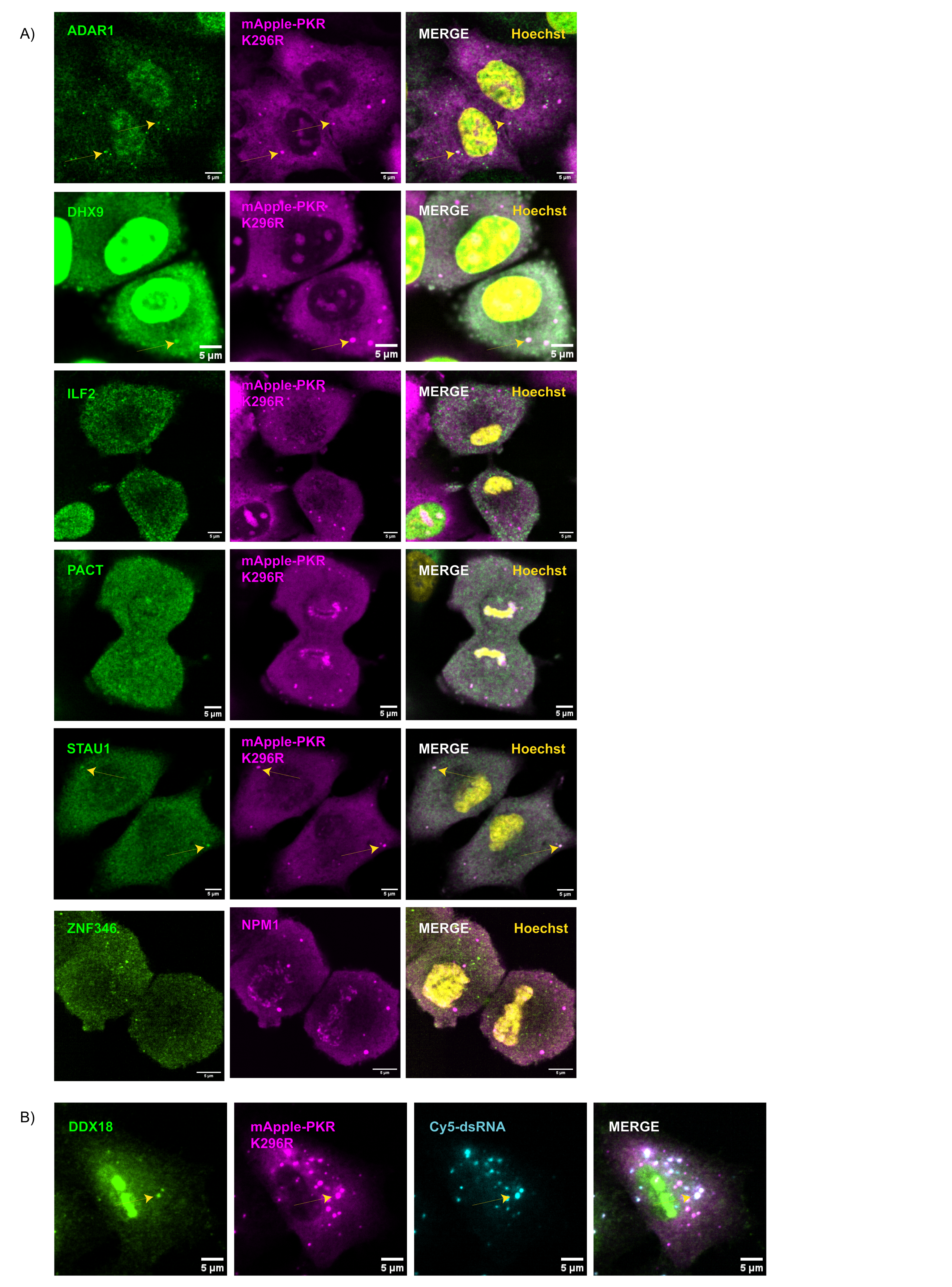


**Figure S4.** (**A**) Representative immunofluorescence microscopy images assessing colocalization between mitotic dRIFs and confirmed $dRIF_{IV}$ resident proteins in A549 PKR KO cells expressing mApple-$PKR^{K296R}$. Image of DHX9 depicts the only instance in which DHX9 proteins was observed in mitotic dRIFs. (**B**) Representative immunofluorescence microscopy image showing DDX18 localization to dRIFs induced by exogenous dsRNA.

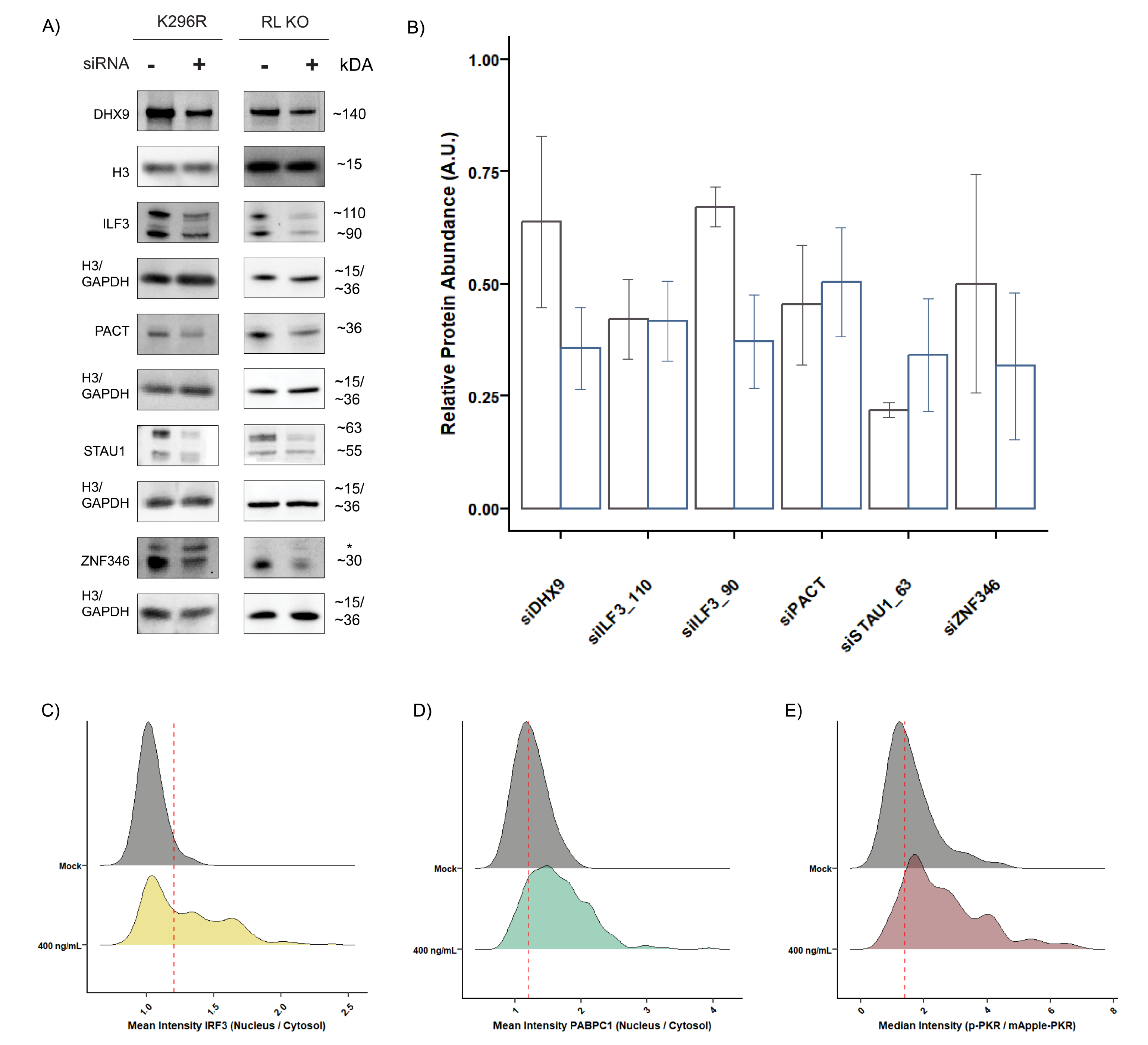


**Figure S5.** (**A**) Representative western blots showing the siRNA knockdown efficiency for each protein target. "*" is used to indicate non-specific bands. (**B**) Quantification of images in S5A and 2-3 additional biological replicates for each condition. Error bars represent the standard deviation of 3-4 independent experiments. (**C-E**) Ridge plots showing the population shift in IRF3 (C) and PABPC1 (D) nuclear versus cytosolic localization in A549 PKR KO mApple-PKR cells, or p-$PKR^{Thr446}$ intensity (E) in PKR/RL dKO mApple-PKR cells upon transfection of 400 ng/mL dsRNA. Vertical line represents the median value of the lipofectamine-only (Mock) negative control condition.

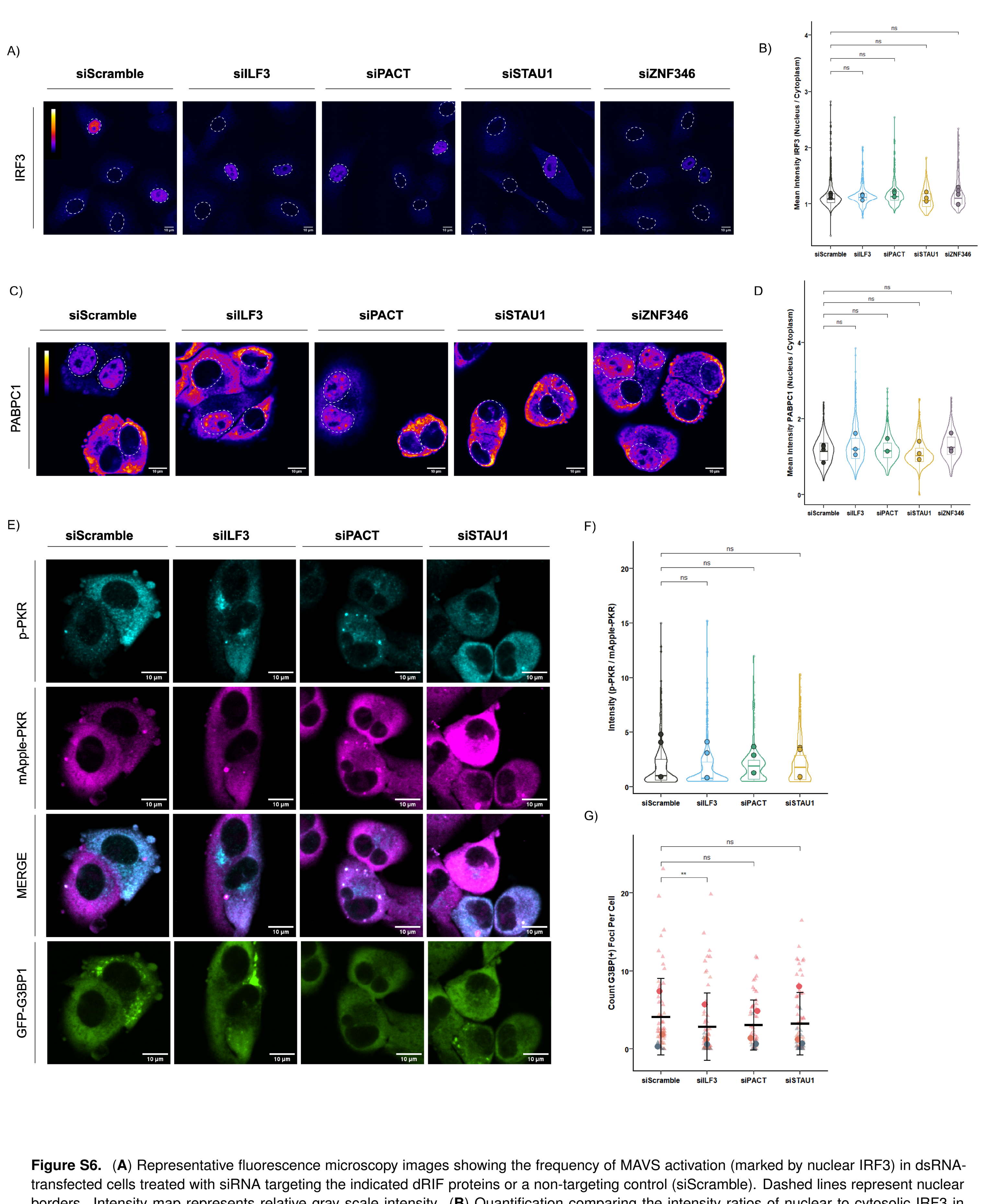


**Figure S6.** (**A**) Representative fluorescence microscopy images showing the frequency of MAVS activation (marked by nuclear IRF3) in dsRNA-transfected cells treated with siRNA targeting the indicated dRIF proteins or a non-targeting control (siScramble). Dashed lines represent nuclear borders. Intensity map represents relative gray scale intensity. (**B**) Quantification comparing the intensity ratios of nuclear to cytosolic IRF3 in dsRNA-transfected cells treated with siRNA targeting the indicated dRIF proteins or a non-targeting control (siScramble). Statistical comparisons were made using a Wilcoxon rank-sum test on per-image means. Data was obtained from 3 independent experiments and 30+ cells and 2+ fields-of-view per biological replicate and condition. (**C**) Representative images showing the frequency of RNase L-activated cells (marked by nuclear PABPC1) in dsRNA-transfected cells treated with siRNA targeting the indicated dRIF proteins or a non-targeting control (siScramble). Dashed lines represent nuclear borders. Intensity map represents relative gray scale intensity. (**D**) Quantification showing the ratio of nuclear to cytosolic PABPC1 intensity in dsRNA-transfected cells treated with siRNA targeting the indicated dRIF proteins or a non-targeting control (siScramble). Statistical comparisons were made using a Wilcoxon rank-sum test on per-image means. Data was obtained from 3-4 independent experiments and 30+ cells and 2+ fields-of-view per biological replicate and condition. (**E**) Representative fluorescence microscopy images showing the relationship between PKR activation, dRIF formation (denoted by mApple-PKR foci) and the presence of stress granules (denoted by GFP-G3BP1 foci) in dsRNA-transfected cells treated with siRNA targeting the indicated dRIF proteins or a non-targeting control (siScramble). (**F**) Quantification showing the mean intensity of p-PKR staining (normalized to mApple-PKR signal) in dsRNA-transfected RL KO cells treated with siRNA targeting the indicated dRIF proteins or a non-targeting control (siScramble). Statistical comparisons were made by performing an unpaired two-tailed t-test on per-image means. Data was obtained from 3 independent experiments and 200+ cells and 2+ fields-of-view per biological replicate and condition. (**G**) Quantification showing the percentage of stress granule-positive cells (proxied by GFP-G3BP1-positive foci) in dsRNA-transfected RL KO cells treated with siRNA targeting the indicated dRIF proteins or a non-targeting control (siScramble). Statistical comparisons were made using an unpaired two-tailed t-test on per-image means. Data was obtained from 3 independent experiments and 100+ cells and 5+ fields-of-view per biological replicate and condition. **All statistical analyses in this figure use the following p-value cutoffs: ****$P \leq 0.0001$, ***$P \leq 0.001$, **$P \leq 0.01$, *$P \leq 0.05$, n.s. $P > 0.05$.

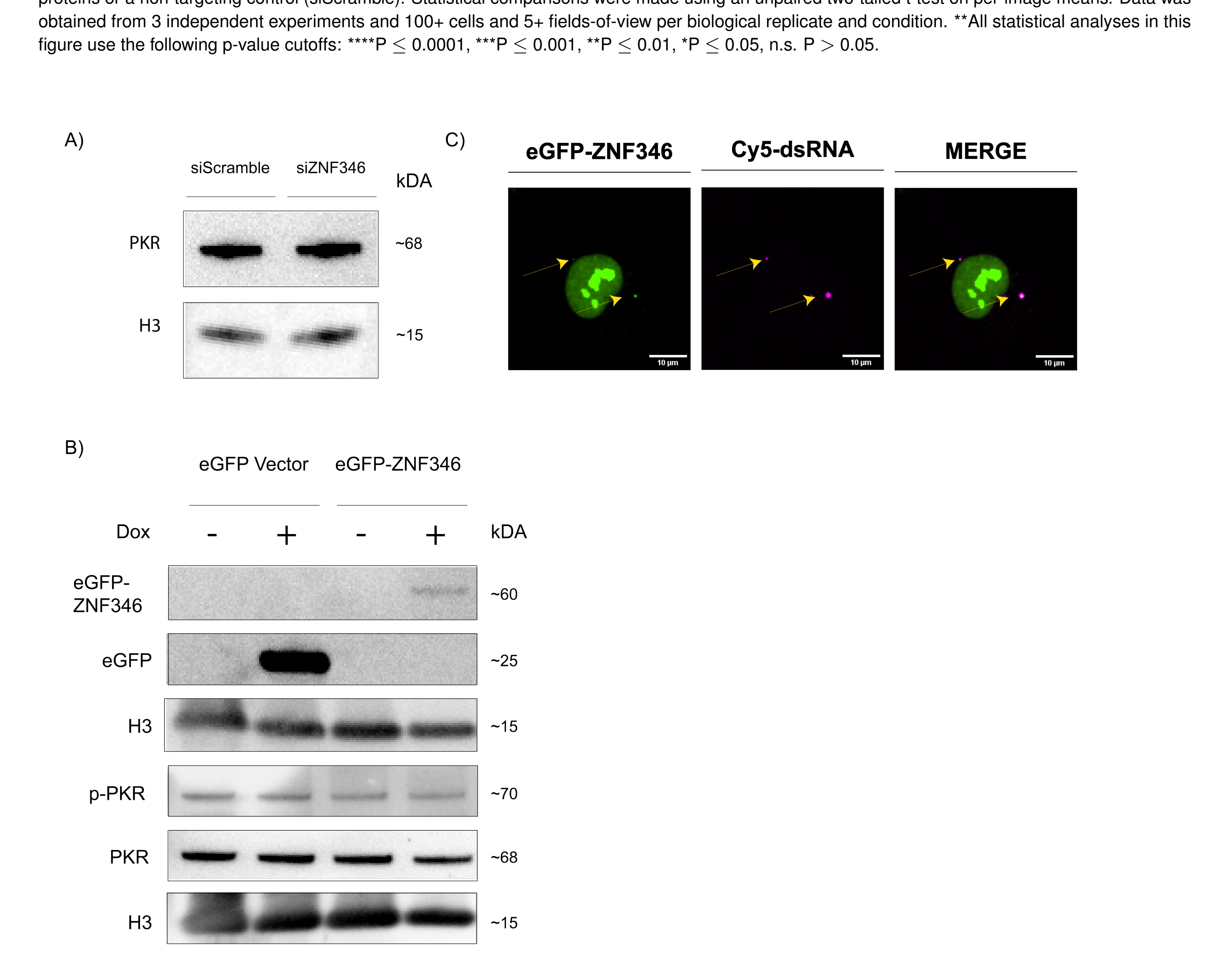


**Figure S7.** (**A**) Western blot of total PKR staining in resting RL KO cells treated with non-targeting control siRNA (siScramble) or siRNA targeting ZNF346 (siZNF346). (**B**) Western blotting for ZNF346, eGFP, PKR, and p-PKR in A549 WT cells expressing either doxycycline-inducible eGFP-ZNF346 or eGFP tagged with V5 with or without 700 ng/mL doxycycline treatment. eGFP-ZNF346 and eGFP bands are cropped for clarity and were imaged on the same membrane at the same exposure time. (**C**) Representative image of an A549 WT cell expressing doxycycline-inducible eGFP-ZNF346 treated with 700 ng/mL doxycycline and 400 ng/mL cy5-labeled dsRNA.
